# Multivalent interactions essential for lentiviral integrase function

**DOI:** 10.1101/2022.01.26.477893

**Authors:** Allison Ballandras-Colas, Vidya Chivukula, Dominika T. Gruszka, Zelin Shan, Parmit K. Singh, Valerie E. Pye, Rebecca K. McLean, Gregory J. Bedwell, Wen Li, Andrea Nans, Nicola J. Cook, Hind J. Fadel, Eric M. Poeschla, David J. Griffiths, Javier Vargas, Ian A. Taylor, Dmitry Lyumkis, Hasan Yardimci, Alan N. Engelman, Peter Cherepanov

## Abstract

A multimer of retroviral integrase (IN) synapses viral DNA ends within a stable intasome nucleoprotein complex for integration into a host cell genome. Reconstitution of the intasome from the maedi-visna virus (MVV), an ovine lentivirus, revealed a large assembly containing sixteen IN subunits (1). Herein, we report cryo-EM structures of the lentiviral intasome prior to engagement of target DNA and following strand transfer, refined at 3.4 and 3.5 Å resolution, respectively. The structures elucidate details of the protein-protein and protein-DNA interfaces involved in lentiviral intasome formation. We show that the homomeric interfaces involved in IN hexadecamer formation and the α-helical configuration of the linker connecting the C-terminal and catalytic core domains are critical for MVV IN strand transfer activity *in vitro* and for virus infectivity. Single-molecule microscopy in conjunction with photobleaching revealed that the MVV intasome can bind a variable number, up to sixteen molecules, of the lentivirus-specific host factor LEDGF/p75. Concordantly, ablation of endogenous LEDGF/p75 resulted in gross redistribution of MVV integration sites in human and ovine cells. Our data confirm the importance of the expanded architecture observed in cryo-EM studies of lentiviral intasomes and suggest that this organization underlies multivalent interactions with chromatin for integration targeting to active genes.

## Introduction

The *Retroviridae* family contains six orthoretroviral genera: alpha-, beta-, gamma-, delta-, epsilon-retroviruses and lentiviruses, as well as a separate subfamily of spumaviruses. Lentiviruses include human immunodeficiency virus type 1 (HIV-1), which is responsible for the global AIDS pandemic. Complementary research into non-human retroviral species greatly accelerated anti-HIV/AIDS drug development from the beginning of the pandemic (2). Retroviral infection proceeds through reverse transcription of the viral RNA genome into a linear double-stranded DNA copy, which is then integrated into a host cell chromosome. Retroviral integrase (IN), the enzyme responsible for this process, catalyzes two consecutive reactions: (*i*) 3’-processing, during which IN hydrolyses 2-3 nucleotides at the viral DNA (vDNA) 3’ ends to liberate 3’-hydroxyls attached to invariant CA di-nucleotides, and (*ii*) strand transfer, wherein IN utilizes the 3’-hydroxyls to cleave the chromosomal DNA, simultaneously joining the 3’ vDNA ends to target DNA (tDNA) strands (reviewed in Ref. (3)). Both reactions proceed via S_N_2 transesterification at phosphorus atoms and require a pair of divalent metal cations (Mg^2+^ or Mn^2+^) as cofactors (4, 5).

To catalyze integration, a multimer of IN assembles into a nucleoprotein complex with synapsed vDNA ends, termed the intasome (6, 7). Results of detailed *in vitro* biochemical and structural studies suggest that the intasome assembles on non-processed vDNA ends as the initial synaptic complex (ISC) that sequentially transitions into the cleaved synaptic complex (CSC, upon 3’-processing of vDNA ends), target capture complex (TCC, upon tDNA binding), and, finally, the post-catalytic strand transfer complex (STC) (5). Disassembly of the STC and subsequent joining of the 5’ vDNA ends to tDNA are thought to depend on cellular machineries. Retroviral INs contain three canonical domains connected by highly divergent linkers.

The amino-terminal domain (NTD) consists of a compact three-helical bundle stabilized by coordination of a Zn^2+^ ion; the catalytic core domain (CCD) features the RNaseH fold and harbors the invariant D,D-35-E catalytic triad; and the carboxy-terminal domain (CTD) adopts an SH3-like beta-barrel structure (reviewed in Ref. (8)). During the last decade, intasomes from five retroviral genera as well as from a yeast retroelement have been structurally characterized in their ISC, CSC, TCC and/or STC forms (1,5,7,9–16). They all share a common functional unit, termed the conserved intasomal core (CIC) assembled around a pair of vDNA ends. The CIC contains two IN CCD dimers, each providing one active site, joined by the exchange of a pair of NTDs. The two halves of the CIC are separated by a pair of CTDs, which act as rigid spacers separating the inner (catalytic) CCDs. The outer CCDs of the CIC do not play a catalytic function. Depending on the viral species, the CIC is decorated by a variable number of IN chains, completing the respective intasome assemblies. Whereas the intasome from the prototype foamy virus (PFV, a spumavirus) comprises a minimal tetrameric IN complex, representing little more in excess of the CIC (7, 9), its lentiviral counterparts can contain as many as sixteen IN chains. Thus, *in vitro* assembly of intasomes from HIV-1 and the closely related red-capped mangabey simian immunodeficiency virus (SIV_rcm_) yielded highly polydisperse populations containing 10-mer, 12-mer, and 16-mer species in various fractions (13,17,18). By contrast, the intasome from maedi-visna virus (MVV, an ovine lentivirus) behaved as a near-homogenous population with a large majority of particles harboring four IN tetramers (1). The smaller HIV-1 and SIV_rcm_ IN-vDNA nucleoprotein complexes could be explained as fragments of the hexadecameric assembly observed in the MVV structure, due to a loss or partial disorder of individual IN subunits (17). These results underscore the utility of MVV intasomes as a model for *in vitro* studies of lentiviral integration.

HIV-1 IN, as well as its counterparts from other lentiviral species, interact with the chromatin-associated host protein LEDGF/p75, which strongly enhances their *in vitro* strand transfer activity (1,19,20). LEDGF/p75 and, to a lesser degree, its paralog HRP2 are largely responsible for the propensity of lentiviruses to integrate into active transcription units (TUs, reviewed in Ref. (21)). While causing variable and usually modest defects in integration efficiency, ablation of LEDGF/p75 in infected cells results in profound changes to the genomic distribution of HIV-1 integration sites (22–26). LEDGF/p75 contains two compact domains connected by a long flexible linker. The N-terminal PWWP domain binds nucleosomes carrying trimethylated histone H3 Lys36 (H3K36me3) (27–29). The IN-binding domain (IBD), which is located close to the C-terminus of the protein (30, 31), binds the IN CCD dimerization interface, with additional salt bridge contacts between the IBD and the IN NTD stabilizing the interaction (32, 33). The modular organization is thought to allow LEDGF/p75 to act as a tether between IN and H3K36me3-containing nucleosomes, which are enriched within TUs and are associated with transcriptional elongation and pre-mRNA splicing activity (34–36).

The biological significance of the expanded intasome architecture observed in lentiviruses remains unclear. If four IN molecules are sufficient to assemble a functional spumaviral (7) or deltaretroviral (14, 15) intasome, why should a lentivirus require many more IN chains to support an equivalent mechanism? Here, we describe high-resolution structures of the MVV intasome in two functional states: (*i*) prior to engagement of tDNA and (*ii*) following strand transfer. The structures reveal fine details of the protein-protein and protein-DNA interactions within the lentiviral intasome, which could not be resolved in the previous cryo-EM reconstructions (1). Using site-directed mutagenesis, we demonstrate that the hexadecameric assembly is essential for MVV IN strand transfer activity and virus infectivity. We also show that the MVV intasome can recruit as many as sixteen LEDGF/p75 molecules, which may allow it to form multivalent interactions with chromatin, potentially allowing it to be more sensitive to the epigenetic status of target chromatin.

## Results

### High-resolution cryo-EM structures of the MVV intasome in two functional states

We collected single-particle cryo-EM data on the MVV intasome assembled in the form of the CSC and purified under conditions used in our previous study (1). The reconstruction was refined to an overall resolution of 3.4 Å (Figs S1a, S2a, Table S1). Although MVV IN strand transfer activity and intasome assembly *in vitro* are strongly stimulated by LEDGF/p75, the host factor dissociates during intasome purification (1). In agreement with the previously reported MVV CSC structure, which was refined to ∼5 Å resolution, the present reconstruction lacked features consistent with LEDGF/p75 (Fig. S2a). To visualize tDNA binding and to potentially enhance host factor retainment, we assembled the STC intasome using a branched DNA construct mimicking a product of strand transfer and purified it in a buffer with reduced salt concentration. We acquired cryo-EM images of the MVV STC intasome particles and refined the resulting reconstruction to an overall resolution of 3.5 Å (Figs S1b, S2a, S3, Table S1). The well-resolved portions of the cryo-EM map encompassed the intasome with 30 base pairs (bp) of tDNA and two copies of the LEDGF/p75 IBD (Figs 1a, 1d, S2a). Because unstructured and flexible regions form the bulk of the LEDGF/p75 molecule (30), it was unsurprising that only the IBD, which directly engages IN, was observed in the map.

**Figure 1.**
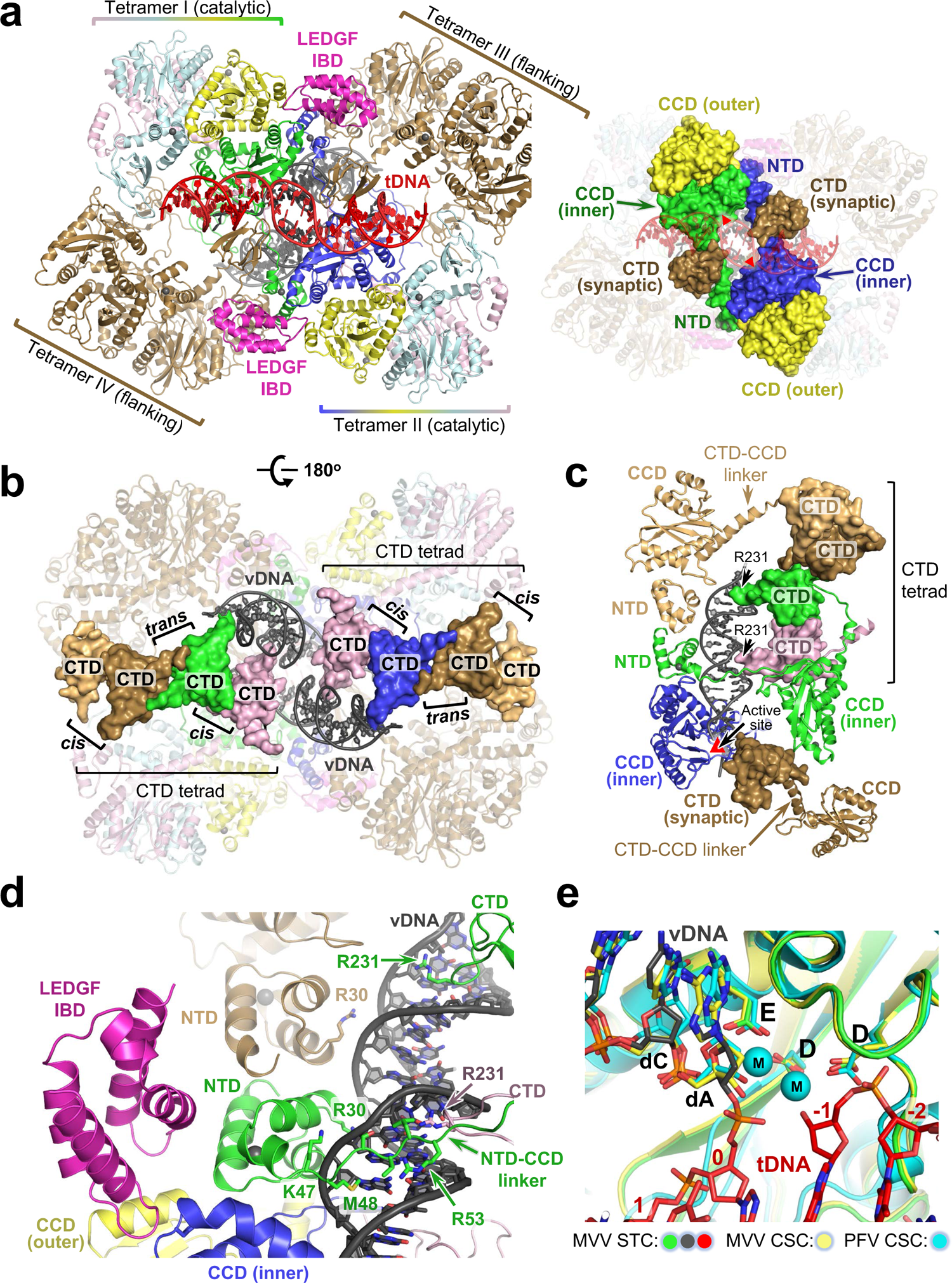
Overview of the MVV intasome architecture. **(a)** Left: Refined model of the STC, shown as cartoons and color-coded to highlight LEDGF/p75 and IN subunits. Catalytic IN tetramers (tetramers I and II) are colored by subunit: the IN chains providing active sites are blue and green, and the remaining chains of the catalytic tetramers are shown in yellow, cyan, and light pink. The eight IN subunits comprising flanking tetramers III and IV are brown. LEDGF/p75 IBDs are magenta; vDNA and tDNA are grey and red, respectively. Right: The STC with IN domains contributing to the CIC shown in surface mode and indicated, and the remainder of the structure shown in semi-transparent cartoons. Red triangles depict active sites. **(b)** The STC with the CTDs comprising the tetrads in surface mode. Interactions between CTDs from same versus different IN tetramers are indicated *cis* and *trans*, respectively. **(c)** IN-vDNA interactions. One of the two vDNA ends is shown. Locations of IN domains (NTD, CCD, and CTD), CCD-CTD linkers, Arg231 residues involved in vDNA binding and the active site (red triangle) are indicated. **(d)** Closeup view of one of the LEDGF/p75 IBDs identified within the STC reconstruction. IN residues involved in the interactions with vDNA are shown as sticks and indicated. **(e)** Closeup view of the MVV STC active site region with IN, vDNA and tDNA in green, grey and red, respectively. Also shown are 3D superimposed structures of MVV CSC (yellow) and PFV CSC (cyan, PDB ID 3OY9 (7)). The three structures were superposed by the Cα atoms of the residues comprising the invariant D,D-35-E motif in each active site (indicated as D, D, and E, corresponding to Asp66, Asp118, and Glu154 in MVV, and Asp128, Asp185, and Glu221 in PFV IN). The protein and DNA are shown as cartoons and sticks, respectively; spheres (M) are catalytic Mn^2+^ cations in the PFV crystal structure (7). Residues of the invariant 3’ vDNA dCdA dinucleotide are indicated, and nucleotides of the tDNA in the MVV STC structure are numbered (0 corresponds to the nucleotide joined to 3’ end of vDNA).

The CSC and STC structures recapitulated the MVV intasome architecture harboring sixteen IN subunits in a tetramer-of-tetramers arrangement. Two IN chains provide active sites for catalytic function, while the remaining subunits are involved in protein-DNA and/or protein-protein interactions. Each of the four IN tetramers making up the intasome contributed to the formation of the CIC, which was resolved to 2.9-3.0 Å resolution within our cryo-EM reconstructions (Figs S2a, S3). The CIC is formed from a pair of CCD dimers provided by IN tetramers I and II, plus a pair of synaptic CTDs derived from IN tetramers III and IV (Fig. 1a). As in all other characterized intasomes, the inner (catalytic) IN chains of the CIC exchange their NTDs across the synaptic interface (3), with the entire extended CCD-NTD linker resolved in the cryo-EM maps (Fig. S3b).

Comparison of the CSC and STC structures confirmed that the intasome does not undergo extensive conformational changes upon tDNA binding and strand transfer (Fig. S2b). In agreement with other retroviral intasome structures (9,12,13,15), the tDNA bound the CIC within the cleft between a pair of catalytic IN CCDs, where it made additional contacts with the synaptic CTDs (Figs 1a, 2a, S3c). Considerable distortion of the duplex structure allowed placement of the tDNA scissile phosphodiester bonds, separated by 20 Å in B-form DNA, into the intasomal active sites spaced by 30 Å. Compared to non-lentiviral species, the MVV intasome induced less extensive tDNA bending into the minor groove, resulting in an obtuse angle between the tDNA arms and a lack of minor groove compression (Fig. S4). MVV achieves the required widening of the tDNA major groove by more extensive stretching and underwinding of the duplex (Fig. S4), resulting in a 4 Å stretch and 22° undertwist of the 6-bp tDNA segment between insertion sites of the vDNA ends. The distortion of the tDNA duplex is stabilized by interactions with IN subunits comprising the CIC. Two pairs of MVV IN Arg231 and His233 residues, located on the synaptic CTDs, along with Trp145 on the inner CCDs of the CIC, make direct contacts within the expanded tDNA major groove (Fig. 2a). Side chains of Arg231 and Trp145 are in proximity with the C5 methyl group of the deoxythymidine at tDNA position −2. The hydrophobic interactions in the major groove may contribute to the strong preference for a thymine base at this position of MVV integration sites and the symmetric preference for adenine at position +7 (Fig. 2b) (1).

**Figure 2.**
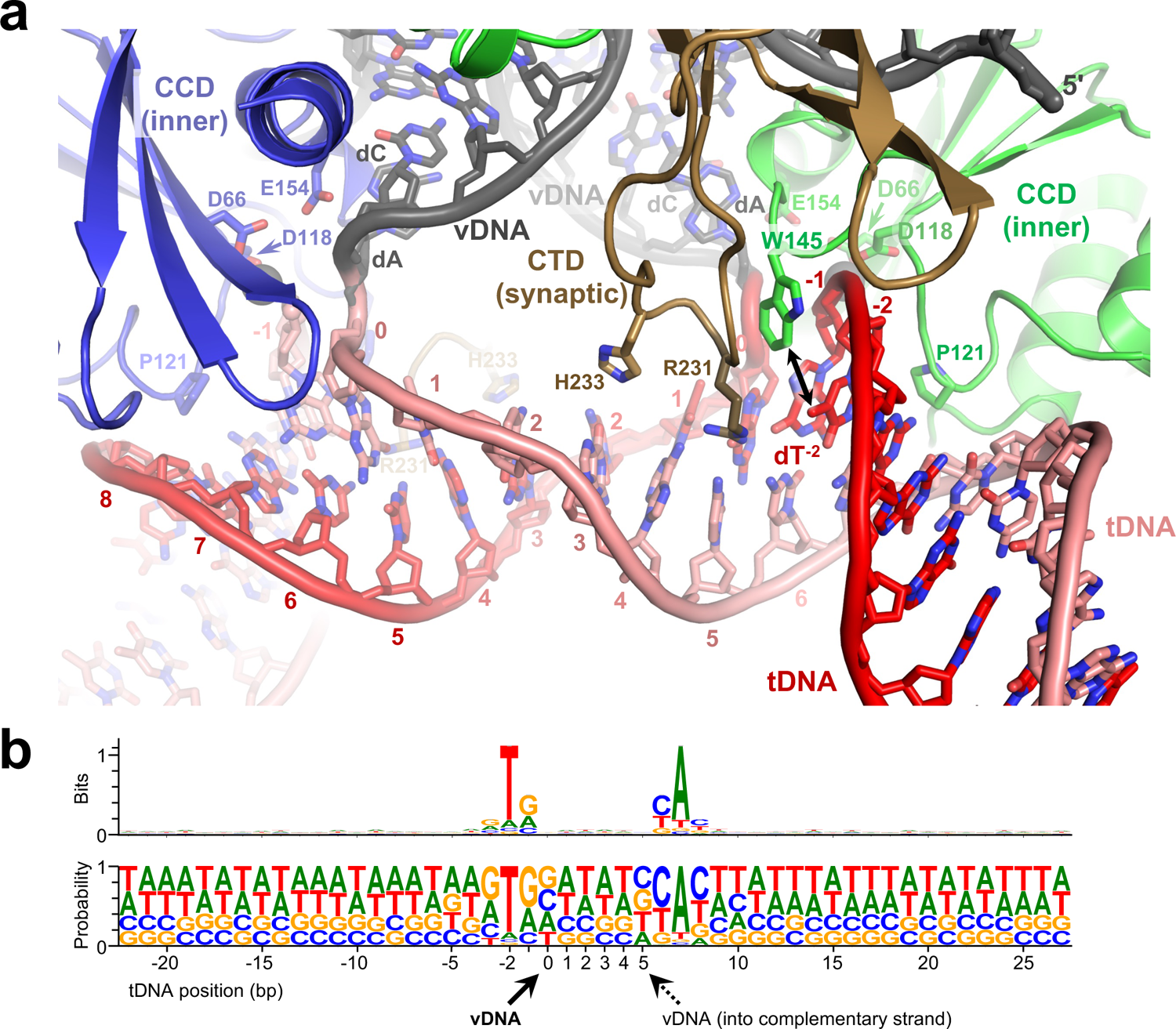
The engagement of tDNA by the MVV intasome. **(a)** Closeup view of the vDNA-tDNA synapse within the MVV STC. DNA phosphate backbone is shown as cartoon with sugar and bases as sticks. Complementary tDNA strands are in red and pink. The rest of the chains are colored as in Fig. 1A. MVV IN residues involved in the interactions with tDNA are shown as sticks and indicated. **(b)** Nucleotide preferences at MVV integration sites. Sequence logos (108, 110) represent information content (reported in bits, top) or raw nucleotide frequencies (bottom) at each position within an alignment of 327,911 *in vitro* MVV intasome integration sites in sheep genomic DNA (1). Nucleotide positions of the tDNA strand represented by the logos are numbered, and the nucleotide that becomes joined to 3’ vDNA end (corresponding to position 0) is indicated with solid black arrowhead. The dotted arrowhead indicates the insertion position of the second vDNA end into the complementary tDNA strand.

During the strand transfer reaction, the tDNA scissile phosphodiester group is coordinated by the catalytic metal ion pair in the IN active site (5). Crystallographic studies of PFV intasomes revealed that strand transfer leads to ejection of the trans-esterified phosphodiester from the active site (5, 9), which is thought to drive the reaction towards integration by discouraging the reversal of transesterification. Although we acquired cryo-EM data in the absence of Mg^2+^ or Mn^2+^ ions, superposition of the structures with the active site region of the PFV intasome, which was crystallized in the presence of Mn^2+^, allowed modelling the catalytic metal ions in the MVV IN active site (Fig. 1e). Within the MVV STC, the phosphodiester linking viral and target DNA molecules is 6 Å apart from the predicted position of either metal ion, indicating that reconfiguration of the active site following strand transfer is conserved in lentiviral intasomes.

### The CTD-CTD interfaces and the α-helical configuration of the CCD-CTD linker are critical for MVV IN strand transfer activity

Retroviral intasomes display striking architectural diversity (3), and outside of the CIC the MVV intasome is distinct from all characterized non-lentiviral intasomes. The cryo-EM structures revealed a critical role played by IN CTDs to stabilize the entire hexadecameric MVV intasome assembly. The CTDs from eight IN subunits form a pair of *C2* symmetry-equivalent tetrads stabilizing the hexadecameric assembly via intra- and inter-tetramer CTD-CTD interactions (Fig. 1b, indicated as *cis* and *trans* tetramer contacts, respectively) as well as via contacts with vDNA (Fig. 1c). CTD dimerization is conserved among lentiviral INs (1,13,17,37), although further multimerization into the tetrad has thus far been observed only for MVV IN in the context of the vDNA-bound intasome (1). Another characteristic feature, unique to lentiviral INs, is the α-helical configuration of the linker connecting the CTD and CCD (1, 38). Importantly, the CTD-CTD interfaces and CCD-CTD linker do not directly contribute to the CIC. Although the CIC contains a pair of CTDs, these assume unique synaptic positions and do not participate in homomeric CTD-CTD interactions.

To test the functional significance of the lentivirus-specific MVV intasome features, we targeted select IN residues by site directed mutagenesis. Amino acid substitutions F223A, Y225A, W245E/L, V252A/D, Y261A/E, V263E, and I272E were designed to disrupt MVV IN intra- and inter-tetramer CTD-CTD interfaces (Fig. 3a, left). To destabilize the CCD-CTD linker (Fig. 3a, right), we produced MVV IN variants harboring substitutions of Gln residues for helix-destabilizing Pro and Gly (QQ207GP and QQQ211PGG). To the same end, we constructed a chimera carrying a portion of the PFV IN CCD-CTD linker (HPSTPPASSRS, corresponding to PFV IN residues 304-314, known to adopt an extended conformation (7)) inserted between MVV IN Gln210 and Gln214, resulting in the variant referred to as QQQ211PFV_304-314_. As controls, we produced MVV IN E154Q, R231E, and H12N variants. Glu154, which is the catalytic glutamic acid of the IN D,D-35-E motif (Fig. 1e), does not contribute to any of the IN-IN or IN-DNA interfaces of the intasome. Therefore, E154Q was expected to abrogate IN catalytic activity, while preserving MVV IN multimerization and intasome assembly. The side chains of Arg231 residues from various IN chains are directly involved in interactions with viral and target DNA (Figs 1c and 2a). Finally, MVV IN His12 is part of the invariant HHCC motif that coordinates a Zn^2+^ ion, which is critical for NTD structural stability (39, 40). Within the MVV intasome, the NTDs from numerous IN chains contribute to the CIC and/or to IN-vDNA and IN-IN interactions (Figs 1a, 1d, and S3b). Therefore, both R231E and H12N were expected to disrupt MVV intasome formation.

**Figure 3.**
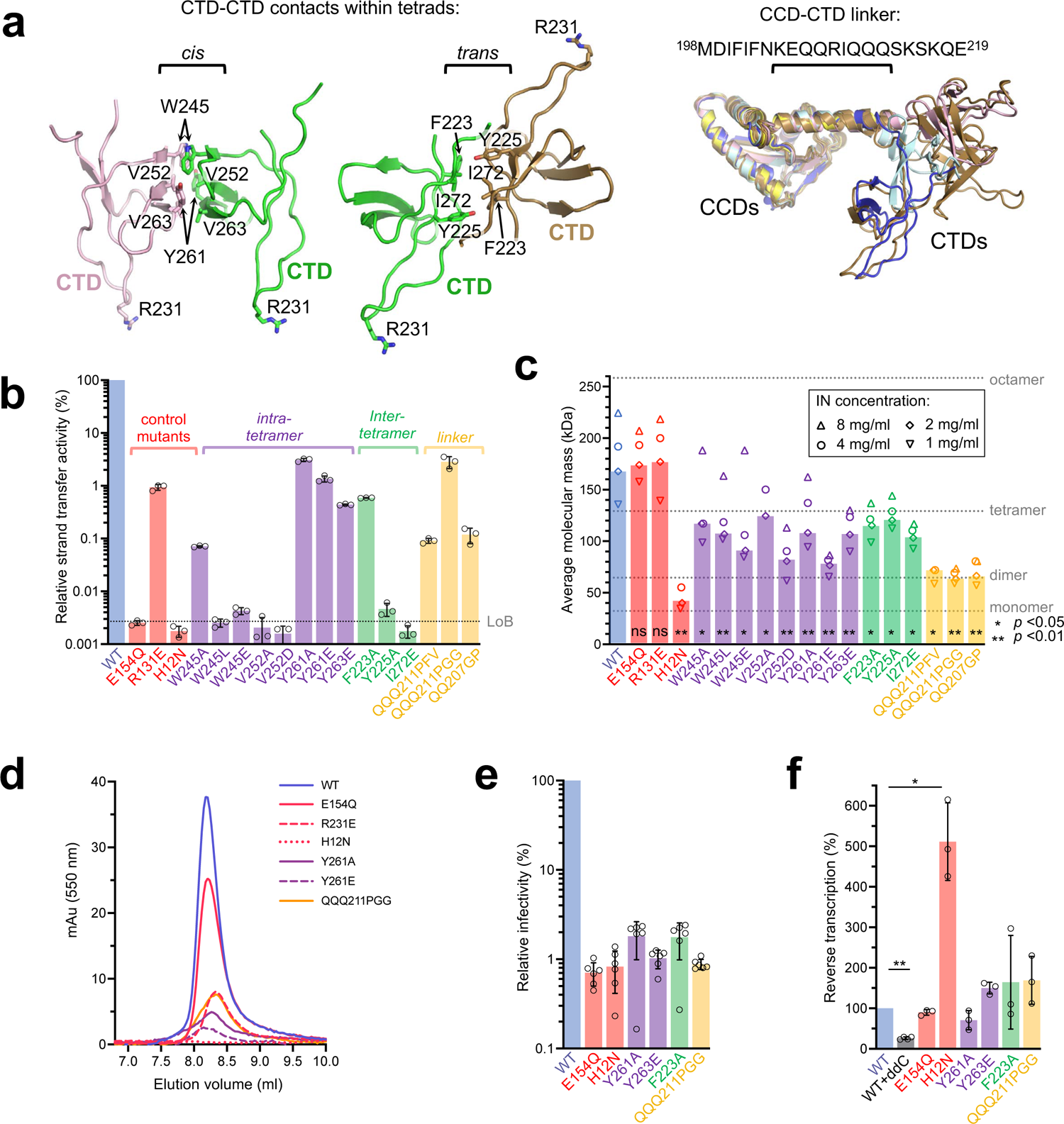
The design and activities of MVV IN mutants. **(a)** Locations of targeted IN residues within intra- and inter-tetramer CTD-CTD interfaces (left) and configuration of the CCD-CTDs linkers within 8 structurally distinct IN chains of the intasome (right; amino acid sequence of the linker is shown above the superposition). Due to *C2* symmetry, the two CTD tetrads (Fig. 1b) are equivalent within the intasome. **(b)** Strand transfer activities of indicated MVV IN mutant proteins relative to that of WT (set to 100%), measured by real-time PCR. The bar plot displays mean values with standard deviations from three replicates for each condition; open circles indicate values for individual repeat measurements. The grey dotted line represents the level of background (LoB), determined from three IN-omit reactions and defined as mean background + 1 standard deviation. Qualitative analysis of the strand transfer products by agarose gel electrophoresis is shown in Fig. S5b. **(c)** Average molar masses (kDa) of MVV IN variants determined by SEC-MALLS upon injections of the proteins at 8, 4, 2, and/or 1 mg/mL (indicated with upward triangles, circles, diamonds, and downward triangles, respectively). Bars represent values obtained with IN injected at 2 mg/mL. Molecular masses of MVV IN monomer (32.3 kDa), dimer, tetramer, and octamer are indicated with grey dotted lines. Statistical significance (WT vs mutant) was estimated using two-tailed paired Student’s t-test, and the results are reported as highly significant (**, *p*<0.01), significant (*, *p*<0.05), or non-significant (n.s.). **(d)** Size exclusion chromatography elution profiles of CSC intasomes assembled with Cy3-labeled vDNA and WT or mutant MVV INs. The curves report Cy3 absorbance at 550 nm to distinguish nucleoprotein complexes from protein aggregates. Only elution volumes 7-10 mL are shown here; the complete elution profiles (0-20 mL), including results of the intasome assemblies with the remaining MVV IN mutants, are shown in Fig. S5d. **(e)** Infectivity of single-cycle MVV-derived vectors produced using Gag-Pol constructs incorporating WT or indicated mutant IN. Luciferase expression was measured 7 d post-infection. The bars indicate mean values with standard deviations from six replicates for each condition; open circles indicate values for individual measurements. **(F)** Quantification of late reverse transcription products in cells infected with WT or IN mutant MVV vectors at 8 h post-infection. The bars show mean values with standard deviations from three biological replicates for each condition; open circles indicate values for individual measurements; two-tailed paired Student’s t-test was used to estimate WT-vs-mutant statistical significance.

We first evaluated the ability of the MVV IN variants to carry out strand transfer *in vitro*, utilizing short double-stranded oligonucleotide mimics of pre-processed MVV U5 vDNA ends. When supercoiled plasmid is used as target, two types of strand transfer products can be separated by agarose gel electrophoresis. Full-site products result from concerted insertions of pairs of vDNA oligonucleotides, linearizing the target plasmid. Uncoupled strand transfer of a single vDNA oligonucleotide yields half-site products, which migrate in agarose gels similar to the open circular form of the plasmid (Fig. S5a). In agreement with published results, WT MVV IN displayed robust strand transfer activity, generating full-site and half-site products in the presence of LEDGF/p75 (Fig. S5b) (1). All MVV IN mutants from our panel displayed profound defects in enzymatic activity, with only F223A, R231E, Y261E, Y261A, V263E, and QQQ211PGG INs generating detectable levels of strand transfer products (Fig. S5b). Quantification of overall strand transfer levels using real-time quantitative PCR revealed 30-fold or greater defects in activity across the mutant panel (Fig. 3b).

### Mutations targeting CTD-CTD interfaces and the CCD-CTD linker perturb MVV IN multimerization and intasome assembly *in vitro*

To study effects of the amino acid substitutions on self-association of MVV IN, we analyzed the proteins by size exclusion chromatography coupled with multi-angle laser light scattering (SEC-MALLS). MVV IN proteins, at 1, 2, 4 and 8 mg/mL, were separated by chromatography through a Superdex-200 column and the molar mass distribution was resolved by MALLS. Under these conditions, WT MVV IN was predominantly tetrameric at the lowest protein concentration and formed larger species, likely dimers of tetramers, at higher input concentrations (Fig. 3c), recapitulating our previous observations (1) as well as similar studies based on HIV-1 IN (41, 42). As expected, E154Q and R231E behaved similarly to WT protein (Fig. 3c). At all input concentrations, WT, E154Q, and R231E INs displayed average molecular masses between those of a tetramer and octamer. The NTD is critical for IN tetramerization (33, 43) and, concordantly, H12N IN eluted as a mixture of monomers and dimers (Fig. 3c). The mutants with substitutions at the CTD-CTD interfaces and the CCD-CTD linker displayed considerable and significant multimerization defects, displaying average molar masses below that of the IN tetramer at input concentrations of 2 and 1 mg/mL (Fig. 3c). This was particularly apparent with the linker variants QQQ211PGG, QQ207GP and QQQ211PFV_304-314_, which were predominantly dimeric, even up to 8 mg/mL.

We reasoned that the reduced ability of MVV IN mutants to assemble into tetramers and/or higher order multimers may affect its ability to form intasomes and thus explain the observed defects in strand transfer activity (Figs 3b, S5b). To test this hypothesis, we monitored MVV IN assembly into intasomes in the presence of vDNA by size exclusion chromatography. As expected, WT protein as well as active site mutant E154Q efficiently formed high-molecular weight nucleoprotein complexes, with elution volumes expected for the intasome (Figs 3d and S5c) (1). While R231E, Y261A, Y261E and QQQ211PGG INs assembled into intasomes with greatly reduced yields (Fig. 3d), the remaining mutants failed to do so (Fig. S5d). Notably, aside from the E154Q active site mutant control, the IN mutants capable of forming detectable intasome complexes also retained detectable levels of strand transfer activity (Fig. 3B).

### Disruption of the IN CTD-CTD interfaces or the α-helical CCD-CTD linker abrogates MVV infectivity

To further investigate the importance of the hexadecameric intasome assembly, the mutations were introduced into a single-cycle MVV virus. We utilized a four-component MVV-derived lentiviral vector system comprising a Gag-Pol packaging construct, a transfer vector harboring the firefly luciferase reporter gene under the control of an internal cytomegalovirus (CMV) promoter, and plasmids expressing MVV Rev and vesicular stomatitis glycoprotein G (VSV-G) proteins (44, 45). We introduced H12N, E154Q, F223A, Y261A, V263E and QQQ211PGG mutations into the IN-coding region of the packaging construct and produced VSV-G-pseudotyped vector particles. Human embryonic kidney 293T (HEK293T) cells were infected with the virus preparations, which were normalized by reverse transcriptase (RT) activity levels. Measurement of luciferase activity 7 d post-infection revealed 50 to 100-fold infectivity defects of the mutants. Notably, infectivities of the F223A, Y261A, V263E and QQQ211PGG viruses were comparable to those of the E154Q or H12N controls (Fig. 3e).

We next evaluated biochemical and virological properties of the IN mutant viruses to assess the nature of the infectivity defects. Immunoblotting of viral lysates revealed that the viruses carrying single IN missense mutations harbored normal levels of mature p24 capsid protein (Fig. S6a). Capsid levels were reduced approximately 3-fold in QQQ211PGG particles. However, this relatively minor maturation defect appeared incongruent with the ∼80-fold defect in infectivity (Fig. 3e). Amino acid substitutions in HIV-1 IN often result in pleiotropic phenotypes, characterized by defects in vDNA synthesis during reverse transcription (46, 47). To determine whether the reduced luciferase levels reflected a defect occurring during integration or at an earlier step of MVV replication, we measured the levels of reverse transcription products in infected cells. As expected, WT and IN mutant reverse transcription in most cases peaked at 8 h post-infection, and WT vDNA synthesis was suppressed by 2’,3’-dideoxycytidine (ddC), a potent inhibitor of MVV RT (48) (Fig. S6b). With the exception of H12N, which unexpectedly displayed a large increase of vDNA accumulation, the mutants did not display drastic perturbations at this stage of the viral life cycle (Figs 3f and S6b). The overcompensation of vDNA synthesis by the H12N IN mutant is in sharp contrast to the vDNA synthesis defects observed with H12N IN HIV-1 (49, 50). While we cannot explain this difference, crucially, the MVV IN mutants targeting the CTD-CTD interactions and the α-helical CCD-CTD linker within the MVV intasome did not display drastic reductions in vDNA levels, consistent with specific defects at the level of intasome assembly and the consequential loss of functional integration.

### Characterization of MVV intasome and LEDGF/p75 stoichiometry

In accordance with previously determined crystal structures (32, 33), the two copies of LEDGF/p75 within our STC reconstruction were bound at the IN CCD dimerization interface, making additional contacts with the IN NTD (Figs 1d, S7a). Harboring an IN hexadecamer, the MVV intasome presents sixteen potential LEDGF/p75 binding sites, all of which appear surface exposed and available for the interaction with the host factor (Fig. S7b). As a result of the two-fold symmetry of the assembly, these sites form eight structurally non-equivalent position pairs. We reasoned that the pair of positions occupied by LEDGF/p75 in our STC structure may have the highest affinity for the host factor. To probe the ability of the MVV intasome to bind additional copies of LEDGF/p75, we used single-molecule total internal reflection fluorescence (TIRF) microscopy. MVV intasomes were assembled using biotin- and Cy3-conjugated vDNA oligonucleotides and LEDGF/p75 labelled with SNAP-Surface649 (Surf649) fluorophore. We purified the intasomes by size-exclusion chromatography to remove free vDNA, captured the nucleoprotein complexes in a streptavidin-coated microfluidic flow cell, and then incubated the immobilized intasomes with LEDGF/p75-Surf649. Individual spots containing Cy3 and Surf649 fluorescent signals were detected, and the decrease of Surf649 fluorescence intensity was monitored during photobleaching under conditions of variable ionic strength (Fig. 4a,b and Movie S1). The results revealed an average of six, four and two LEDGF/p75 subunits bound per MVV intasome in the presence of 0.2, 0.5 and 1 M NaCl, respectively, with up to sixteen LEDGF/p75 subunits bound per intasome at the lowest NaCl concentration (Fig. 4c and Table S2).

**Figure 4.**
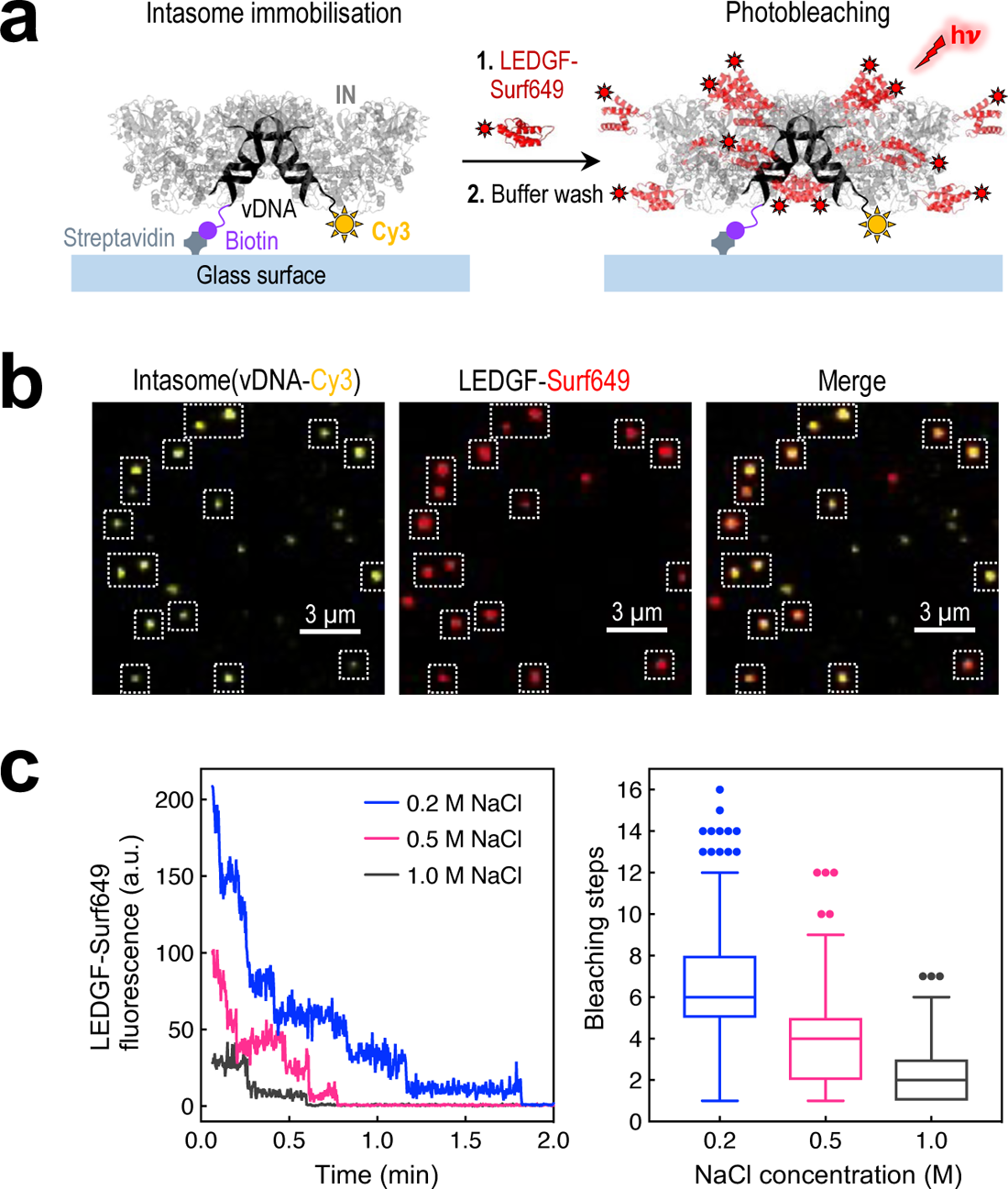
Quantitation of intasome-LEDGF/p75 stoichiometry by single-molecule TIRF microscopy. **(a)** Schematic of the photobleaching experiment. Intasomes containing biotinylated and Cy3-labeled vDNA were immobilized on streptavidin-coated glass cover slips. Following incubation with Surf649-labeled LEDGF/p75 and a wash with buffer supplemented with 0.2 - 1 M NaCl, intasomes were observed by TIRF microscopy enabling the individual steps of Surf649 photobleaching during illumination with a 640 nm laser (hν) to be counted. **(b)** Representative images of surface attached intasome (vDNA-Cy3, yellow; LEDGF/p75-Surf649, red) molecules. Dotted, white line squares indicate individual intasome-LEDGF/p75 complexes. **(c)** Left: Examples of stepwise photobleaching traces of LEDGF/p75-Surf649 at increasing NaCl concentration: 0.2 M (blue), 0.5 M (magenta) and 1.0 M (dark grey). The vertical axis represents fluorescence in arbitrary units (a.u.). Right: Tukey (box-and-whiskers) plots summarizing statistical analysis of the number of LEDGF/p75-Surf649 photobleaching steps per intasome, for various NaCl concentrations (see Table S2 for further details). Each box encloses 50% of the data with the median value displayed as a horizontal line. Outliers are indicated with closed circles.

### LEDGF/p75 strongly influences MVV integration site selection

LEDGF/p75 strongly stimulates *in vitro* strand transfer activity of MVV IN and other lentiviral INs (1,17,19,20). To test if the presence of LEDGF/p75 or its paralog HRP2 (30) in target cells influences MVV infectivity, we used HEK293T-derived cell clones ablated for one or both of these host factors. LKO is an LEDGF-null cell line, generated via TALEN-mediated *PSIP1* gene disruption (51). In addition, we established a dual knockout clone, LHKO, which additionally lacked HRP2 due to disruption of *HDGFL2* gene (Fig. S8a). As above, we infected these cells with RT-normalized quantities of WT or IN active site mutant E154Q MVV vectors. Measured 7 d post-infection, the infectivity of the WT vector in LKO and LHKO cells was reduced ∼5-fold compared to that in parental HEK293T cells (Fig. S8a). The infectivity of the E154Q IN active site mutant virus in HEK293T cells was ∼2% of the WT vector, with the residual luciferase expression likely explained by persistence of non-integrated vDNA forms. However, WT MVV vector infectivity was not restored upon ectopic expression of ovine LEDGF/p75 in LHKO cells (Fig. S8b). Hence, the lack of the host factor seemed unlikely to explain the observed reductions in MVV infection in these cells; instead, the clonal nature of LKO and LHKO cells may account for the reduced transduction efficiency. We accordingly next established a panel of ovine cell lines depleted of HRP2 and/or LEDGF/p75 via synthetic RNA-directed CRISPR-Cas9 genome modification (Fig. S9a). The parental cell line, CPT3, was derived from sheep choroid plexus cells immortalized by co-expression of simian virus 40 large T antigen and human telomerase (52). Importantly, the resulting knockout cell lines (CPT3-LKO1, 2, 3, and 4, and CPT3-LHKO1 and 2) did not undergo single cell cloning. Transduction of the knockout cell panel with the MVV vector did not reveal consistent infectivity defects associated with LEDGF/HRP2 knockout status (Fig. S9b), indicating that the host factors are not essential for MVV infection.

LEDGF/p75 and, to a lesser degree, its paralog HRP2 direct HIV-1 integration towards bodies of highly expressed genes (22,25,26,35,53). To test if LEDGF/p75 plays a similar role in guiding MVV integration, we mapped MVV vector integration sites in our panel of knockout cell lines. Human HEK293T, LKO, LHKO cells and ovine CPT3, CPT3-LKO1, CPT3-LHKO1, and CPT3-LHKO2 cells were infected with the WT MVV vector, and genomic DNA was isolated 5 d post-infection. Chromosomal junctions at integrated U5 vDNA ends were amplified using linker-mediated PCR, sequenced using Illumina technology and aligned with human and sheep genomes. We then correlated the distributions of uniquely mapped MVV integration sites with positions of annotated genes and their transcriptional activity, locations of transcription start sites (TSSs), CpG islands as well as with local A/T content and gene density (Table 1 and 2). In addition, as the locations of human constitutive lamina-associated regions (cLADs, corresponding to perinuclear heterochromatin) and nuclear speckle-associated domains (SPADs, corresponding to transcriptionally-active regions) were available (54, 55), these features were also used in the analyses of MVV integration sites in human cells (Table 1).

**Table 1.**
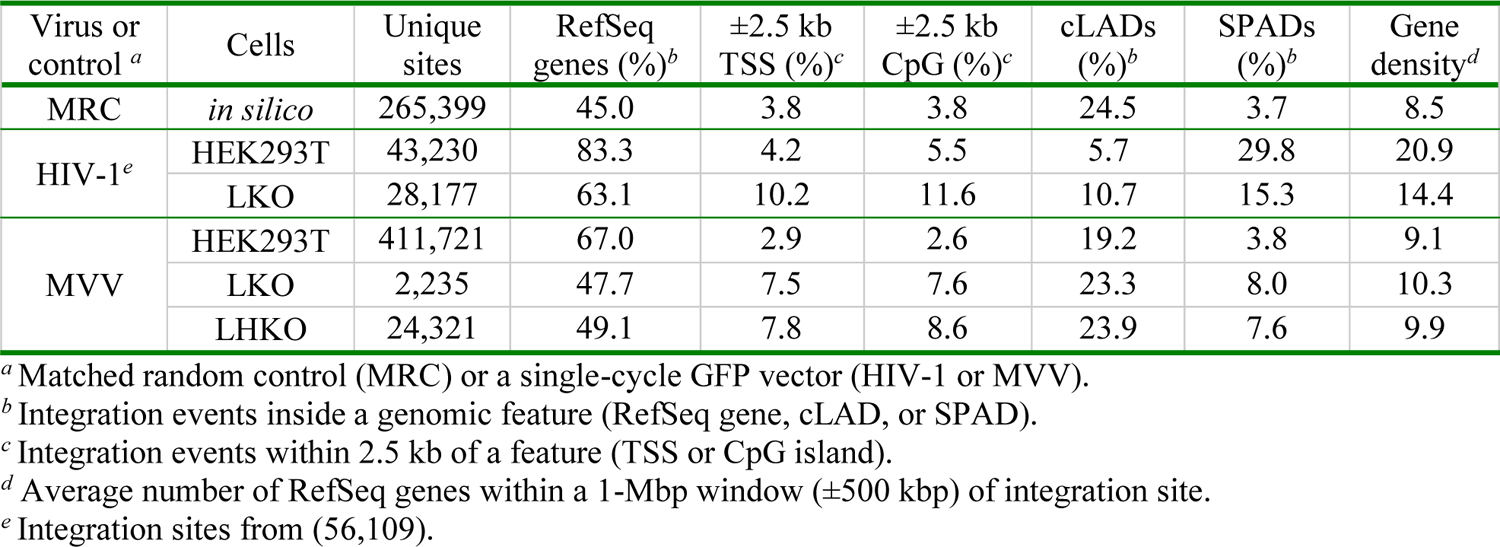
HIV-1 and MVV integration site distributions in human HEK293T, LKO and LHKO cells.

These analyses revealed that MVV displayed a strong bias towards integration into gene bodies in both human (HEK293T) and ovine (CPT3) cell lines (Tables 1 and 2; results of statistical tests are given Tables in S3 and S4). In total, 72.1% of MVV integration sites were found within annotated transcription units in the sheep genome, compared to the expected frequency of 43.3% determined using *in silico*-generated matched random control (MRC). Similar to HIV-1 and other lentiviruses (22,25,56), the frequency of MVV integration into genes significantly decreased in LEDGF-null human (*p* ∼ 10^-77^) and ovine (*p* < 10^-300^) cells, although we did not reproducibly observe an additional defect upon HRP2 ablation. The frequency of MVV genic integration events strongly correlated with transcription activity. Thus, genes expressed at the highest level were ∼4-fold more likely to host an MVV integration event than those expressed at the lowest level; this correlation degreased significantly in the absence of LEDGF/p75 in both human (Fig. 5a) and ovine (Fig. 5b) cells. Similar to HIV-1, the frequency of MVV integration in proximity of TSSs and CpG islands significantly increased in LEDGF-null cells. LEDGF/p75 contains an AT-hook, which was implicated in DNA binding (29, 57). Concordantly, LEDGF/p75 depletion shifted HIV-1 (22, 25) (Fig. 5c) and MVV (Fig. 5c,d) integration events towards regions with lower A/T content. In accordance with the cryo-EM structure (Fig. 2a), ablation of HRP2 and/or LEDGF/p75 did not affect local nucleotide preferences at the integration sites (Fig. S9c), which remained basically unchanged from those observed with *in vitro*-assembled MVV intasomes (Fig. 2b). By sharp contrast to HIV-1, MVV did not display a strong preference for gene-dense regions and SPADs, and only moderately avoided integration into cLAD perinuclear chromatin. In HEK293T cells, HIV-1 integrated into SPADs ∼8-fold more frequently than expected, and this preference was reduced ∼2-fold upon LEDGF/p75 ablation (Table 1). By contrast, the frequency of MVV integration into SPADs nearly precisely matched the expected value of 3.7% and increased 2-fold in LKO (*p* ∼ 4⋅10^-19^) and LHKO cells (*p* ∼ 10^-149^). Although we have not studied the effect of HRP2 ablation in isolation, distributions of MVV integration sites in double knockout LHKO and CPT3-LHKO cells tended to mirror those in LKO and CPT3-LKO, respectively.

**Figure 5.**
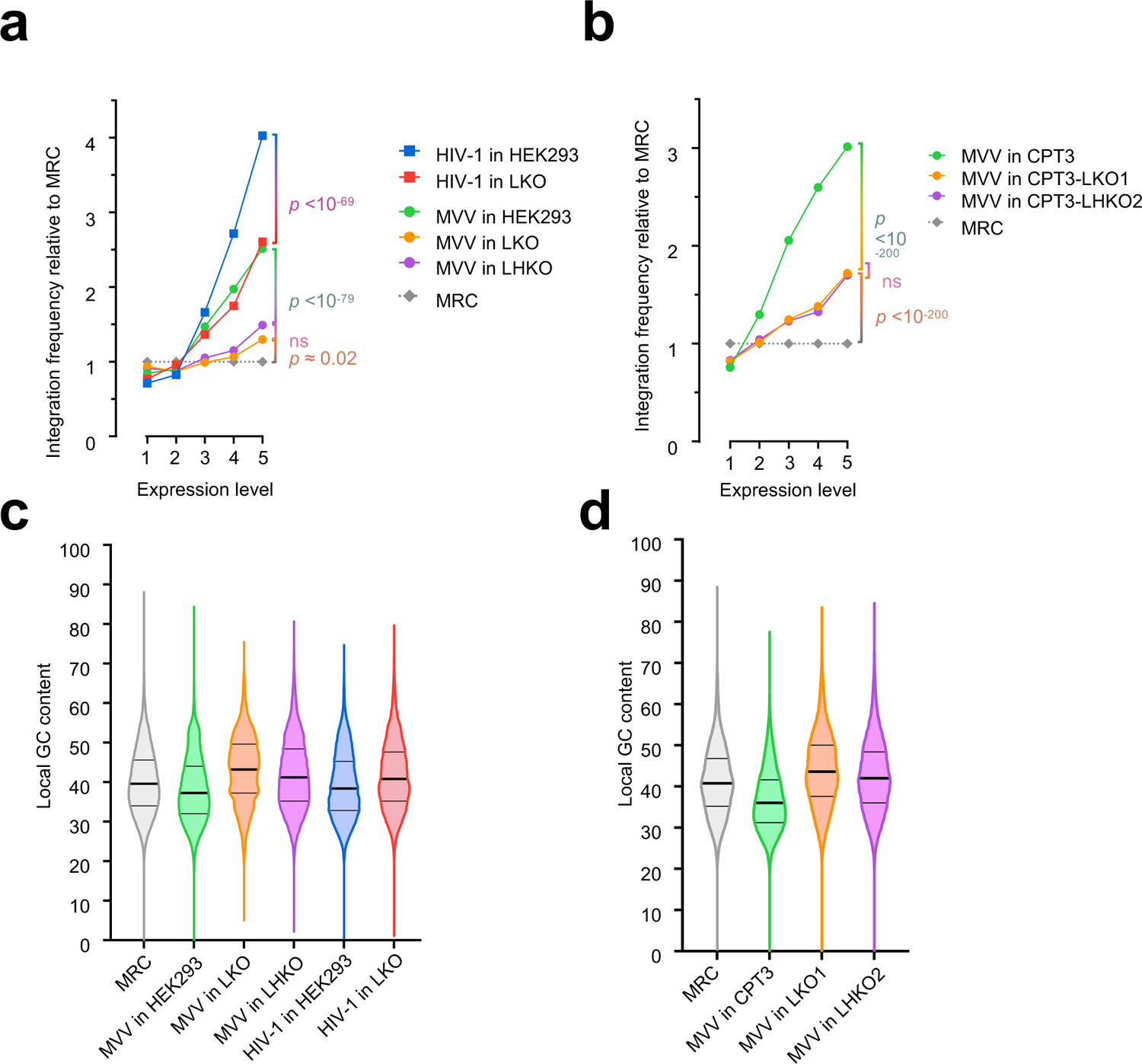
Effects of LEDGF/p75 depletion on MVV integration site distribution. **(A)** Frequency of MVV and HIV-1 integration events into TUs of variable transcriptional activity in HEK293T, LKO, and LHKO cells. Only integration events mapped to RefSeq genes were considered for this analysis. TUs were ranked by their activity into five bins, where each bin contained the same fraction of the genome. Statistical significance (HEK293T vs LKO or LHKO) was determined using Chi-squared test. **(B)** Frequency of MVV integration into TUs of different activity in ovine CPT3, CPT3-LKO1 and -LHKO2 cells. **(C, D)** Local GC contents for mapped integration sites in human (**C**) and ovine (**D**) cells. The data are plotted as violin plots showing frequency distribution for individual GC contents. Thick horizontal lines represent median values, while thin lines indicate boundaries of 25^th^ and 75^th^ percentiles of data points. See Table S3 for statistical analyses of panel C and D datasets.

**Table 2.** MVV integration site distributions in ovine CPT3 cells in the presence and absence of HRP2 and/or LEDGF/p75.

| Virus or control<sup>a</sup> | Cells | Unique sites | RefSeq genes (%)<sup>b</sup> | ±2.5 kb TSS (%)<sup>c</sup> | ±2.5 kb CpG (%)<sup>c</sup> | Gene density |
| --- | --- | --- | --- | --- | --- | --- |
| MRC | *in silico* | 284,737 | 43.3 | 4.6 | 6.1 | 19.6 |
| MVV | CPT3 | 296,801 | 72.1 | 5.2 | 4.4 | 26.8 |
|  | CPT3-LKO1 | 145,863 | 51.8 | 12.4 | 16.5 | 29.1 |
|  | CPT3-LHKO1 | 3,880 | 47.6 | 10.9 | 14.9 | 27.9 |
|  | CPT3-LHKO2 | 82,570 | 51.6 | 12.4 | 16.0 | 28.2 |

<sup>a</sup> Matched random control (MRC) or a single-cycle MVV GFP vector.
<sup>b</sup> Integration events inside a genomic feature (RefSeq gene, intergenic region).
<sup>c</sup> Integration events within 2.5 kb of a feature (TSS or CpG island).

## Discussion

Our cryo-EM reconstructions provide snapshots of a lentiviral intasome in its two key functional states: prior to capture of tDNA and following strand transfer. In agreement with the previous low-resolution structure (1), the MVV intasome contains sixteen IN subunits in a tetramer-of-tetramers arrangement. By comparison, *in vitro* assembly of HIV-1 and SIV_rcm_ intasomes led to formation of highly heterogenous nucleoprotein complexes containing from four to sixteen IN subunits, with a prominence of dodecamers, as well as linear stacks of the intasomes (13,17,18). These observations underscore the utility of the MVV intasome as a convenient model to study lentiviral integration and call for further systematic studies of the lentiviral intasome architecture and its functional significance.

Notwithstanding their heterogeneity, the primate lentiviral intasomes are closely related to their MVV counterpart (see Ref. (17) for details). Importantly, the lentiviral intasomes share two unique features: intra-tetramer CTD-CTD contacts and an α-helical configuration of the CCD-CTD linkers (Fig. 3a). These features do not participate in the formation of the CIC, which is the functional core of the retroviral intasome. The MVV intasome also employs intra-tetramer CTD-CTD interactions within CTD tetrads (Figs 1b and 3a). In solution, MVV IN exists as tetramers, which can self-associate into larger multimers (Fig. 3c) (1). Predictably, disruption of the CTD-CTD or the CCD-CTD α-helical linker significantly reduced the multimerization state of the protein (Fig. 3c). Moreover, the mutations rendered MVV IN unable to catalyze strand transfer, assemble into intasomes *in vitro* and caused gross defects in the context of a single-round infection (Fig. 3). These results argue that the MVV IN evolved to function in a highly multimeric form and is non-functional when forced into smaller sub-complexes. Because HIV-1 particles package ∼100 Gag-Pol molecules, the number of available IN subunits is unlikely to limit intasome assembly during lentivirus infection (58, 59). Moreover, given the proclivity of lentiviral INs to multimerize (Fig. 3c) (19,33,60), we cannot rule out that lentiviral pre-integration complexes may contain even larger IN assemblies. While ascertaining the true multimeric state of lentiviral IN during integration may require visualization of viral nucleoprotein complexes in infected cells, our results argue that the functional intasome must assemble from multiple IN tetramers.

A common property of lentiviral INs is the interaction with LEDGF/p75 (20, 23), which was shown to direct HIV-1 integration towards active TUs (22, 25). Similarly, LEDGF/p75 regulates integration of non-primate lentiviruses such as feline immunodeficiency and equine infectious anemia viruses (56, 61), and, as we have shown here, plays the dominant role in directing MVV integration (Fig. 5). We propose that the ability of the lentiviral intasome to bind a large number of LEDGF/p75 molecules (Figs 4c, S7b) allows the PIC to establish highly multivalent interactions with chromatin. Multiple simultaneous contacts with H3K36Me3-containing nucleosomes can only form in locations enriched for the epigenetic mark. Such a mechanism may allow lentiviral PICs to discriminate H3K36Me3 peaks on the backdrop of noisy epigenetic landscapes, providing a possible biological rationale for the expansion of the lentiviral intasome. Interestingly, similar to other non-primate lentiviruses (56), MVV does not show preference for integration into SPADs (Table 1). This observation is consistent with the idea that recruitment of CPSF6, the factor responsible for HIV-1 targeting to nuclear speckles, is a recent acquisition, specific to primate lentiviruses (56).

Notwithstanding the profound effects on lentiviral integration site distribution, ablation of LEDGF/p75 results in highly variable and often modest defects in single-round HIV-1 infection (Fig. S9b) (22-25,56,61,62). Similarly, redirection of gamma-retroviral integration via mutations disrupting IN binding to cellular bromodomain and extra-terminal proteins revealed little effect on viral infectivity (63). Even more strikingly, despite redirecting a considerable fraction of proviruses to perinuclear heterochromatin, ablation of CPSF6 slightly increases infectivity of HIV-1 and other primate lentiviruses in single-cycle assays (56). Modest infectivity defects were also observed in spumaretroviruses where the Gag mutant R540Q, quantitatively redirected PFV integration into centrosomes (64). Therefore, given these observations, we conclude that single-cycle infectivity assays do not replicate the conditions that led to the emergence of the observed integration preference traits. Integration site selection is likely to confer subtle transcriptional advantages that nonetheless translate to important growth advantages within the infected host, *e.g*., to outpace virus-induced cytopathicity and/or humoral immune pressure (65).

Our structures highlight general features of the retroviral integration machinery and reveal intriguing differences between viral genera. Notably, as in the case of PFV, MVV strand transfer appears to result in ejection of the phosphodiester bond linking viral and target DNA from the IN active site, suggesting conservation of the mechanism proposed to discourage the reversal of the integration process (5, 9). Indeed, a much more distantly related Mu phage transposase was also shown to utilize this mechanism (66). The ejection of the transesterified phosphodiester group is likely driven by tension caused by sharp deformation of the tDNA duplex. The size of the tDNA segment separating insertion sites of the two vDNA ends into opposing tDNA strands ranges between 4 and 6 bp, depending on the retroviral species. By contrast, the distance between intasomal active sites remains relatively constant between retroviral intasomes (28-30 Å, defined as the distance between Cα atoms of the invariant IN active site Glu residues). This explains the marked differences in tDNA conformation induced by diverse retroviral intasomes (Fig. S4). Evidently, in each case, the observed deformation allows sufficient expansion of the tDNA major groove to afford intasomal active sites access to the scissile phosphodiester bonds. The propensity of retroviruses and retrotransposons to integrate into nucleosomes is supported by strong evidence (67–69), and the PFV intasome in complex with a core nucleosome particle was visualized by cryo-EM (70, 71). However, wrapping around the histone octamer imposes constraints of the conformation and major groove accessibility of nucleosomal DNA (72). Thus, the differences in tDNA conformation during its capture by the retroviral intasomes strongly support the idea that retroviruses may have evolved genus or even species-specific ways of interacting with chromatinized target DNA (73).

## Materials and Methods

### Recombinant protein expression and purification

The plasmid pCPH6P-MVV-IN, encoding IN from the KV1772 MVV isolate (74) with a cleavable hexahistidine tag, was previously described (1). Mutations were introduced into pCPH6P-MVV-IN using PCR. Plasmid pET22b-MVV-CAfl, encoding the MVV capsid/p24 protein, was constructed by Gibson assembly as recommended by the manufacturer (New England Biolabs) using PCR fragment amplified from pCAG-MV-GagPol-IN^KV1772^-CTEx2 (44, 45) and *Nde*I/*Xho*I-linearized pET-22b(+) (Millipore Sigma). MVV IN proteins and human LEDGF/p75 were produced in bacteria and purified as previously described (1). The construct used for expression of LEGDF-SNAPf, which was used in TIRF microscopy, was made by ligating a PCR fragment encoding full-length human protein between *Apa*I and *Bam*HI sites of pHis_10_-PS-SNAPf (75) (gift from Ron Vale; Addgene plasmid #78512; http://n2t.net/addgene:78512; RRID:Addgene_78512). LEGDF-SNAPf protein was labelled with SNAP-Surface649 (Surf649, New England Biolabs). Briefly, 8 μM LEDGF/p75-SNAPf was incubated with 16 μM Surf649 in 500 mM NaCl, 5 mM DTT, 50 mM Tris-HCl, pH 7.5 for 30 min at 37°C. Labelled protein was purified from excess dye using a Bio-Spin P6 column (BioRad); labelling was monitored by native protein mass spectrometry (Mass Spectrometry Facility, University of St. Andrews, UK).

Recombinant MVV capsid/p24 protein was produced in *E. coli* BL21(DE3) cells transformed with pET22b-MVV-CAfl. Cells were grown in shaker flasks at 37 °C to an OD_600_ of 0.6, and protein expression was induced with 1 mM isopropyl β-D-1-thiogalactopyranoside for 4 h at 37 °C. Pelleted cells resuspended in lysis buffer (50 mM Tris-Cl pH 8.0, 50 mM NaCl, 20 mM imidazole, 1 mM TCEP) were flash-frozen. Thawed resuspensions were lysed by sonication for 2 min on ice in 5 s intervals, with 30 s gaps between bursts. The supernatant fraction following centrifugation at 50,000*g* for 30 min was loaded onto a HisTrap HP column (GE Healthcare). After extensive washing with lysis buffer, the column was developed with elution buffer (50 mM Tris-Cl pH 8.0, 50 mM NaCl, 250 mM imidazole, 1 mM TCEP). Eluted protein, concentrated by ultrafiltration, was loaded onto a Superdex S200 10/300 gel-filtration column equilibrated in SEC buffer (50 mM Tris-Cl pH 8.0, 50 mM NaCl, 1 mM TCEP). Pooled fractions of capsid protein were flash-frozen in liquid nitrogen and stored at −80°C. For antibody production, thawed protein was dialyzed against phosphate-buffered saline (PBS) overnight at 4 °C. The protein was used for production and affinity purification of polyclonal rabbit anti-CA/p24 antibody (Thermo Fischer Scientific).

### MVV intasome assembly and purification

HPLC-purified oligonucleotides were purchased from Sigma-Aldrich. The MVV CSC intasome complex was assembled using a double-stranded oligonucleotide mimicking the terminal 29 bp of the processed MVV U5 vDNA end prepared by annealing synthetic oligonucleotides EV272 (5’-CCGTGCAACACCGGAGCGGATCTCGCA) and EV273 (5’-GCTGCGAGATCCGCTCCGGTGTTGCACGG). When required, EV272 was modified with 5’-Cy3 dye (for TIRF microscopy, photobleaching and quantitative intasome assembly assays) and EV273 with 3’-triethylene glycol (TEG) biotin (in TIRF microscopy and photobleaching experiments). The MVV STC was assembled using a DNA construct corresponding to the product of full-site integration of 23-bp MVV U5 viral DNA end (vDNA) mimic into a palindromic 52-bp target DNA. The branched DNA was made by annealing oligonucleotides 5’-GCTGCGAGATCCGCTCCGGTGTT, 5’-AACACCGGAGCGGATCTCGCA<u>GTCGACCACCCTAATCAAGTTTTTTGGGG</u> and 5’-<u>CCCCAAAAAACTTGATTAGGGTG</u>; the palindromic tDNA portion (underlined) was designed based on the preferred integration site sequence in pGEM9zf plasmid (1). The intasomes were typically assembled by incubating 7 μM MVV IN, 8 μM LEDGF/p75, and 3.75 μM annealed vDNA (or the strand transfer product mimic) in 80 mM NaCl, 40 mM potassium acetate, 3 mM CaCl_2_, 10 μM ZnCl_2_, 1 mM dithiothreitol (DTT) and 25 mM BisTris-HCl, pH 6.0 in a total volume of 200 μl at 37 °C for 10 min. To prepare samples for cryo-EM, the reaction was upscaled to a total volume of 1 mL. The opalescent mixture was supplemented with 50 mM BisTris-HCl, pH 6.5 and 190 mM NaCl and incubated on ice for 5 min to clear. If the starting volume exceeded 200 μL, the mixture was concentrated by ultrafiltration in a VivaSpin device to a final volume of 200 μL. Intasomes were purified by size exclusion chromatography on a Superdex-200 10/30 column (GE Healthcare) pre-equilibrated in 310 mM NaCl, 3 mM CaCl_2_, 25 mM BisTris-HCl, pH 6.5.

### MVV CSC cryo-EM grid preparation, data collection and image processing

MVV CSC intasomes, assembled and purified as previously described (1), were applied onto R1.2/1.3 gold UltrAufoil grids, Au 300 mesh (Quantifoil). Cryo-EM grids were prepared by manually freezing using a manual plunger in cold room at 4°C and stored in liquid nitrogen for future data acquisition. Cryo-EM movie frames were collected on a Talos Arctica transmission electron microscope (Thermo Fisher Scientific) operating at 200 keV equipped with a K2 summit direct detector (Gatan). Data collection was performed using the Leginon software (76, 77) at a magnification of 45,000, corresponding to a pixel size of 0.92 Å/pixel in nanoprobe mode. The stage was tilted to 40° during data collection to account for the preferential orientation of the sample within the vitreous ice (78). Movies composed of 100 frames were collected in counting mode over 10 s (100 ms per frame). The total fluence was 43.6 e^-^/Å^2^ at a rate of 3.7 e^-^/pix/sec. Imaging parameters are summarized in Table S1.

The movie frames were motion-corrected and dose-weighted using MotionCor2 (79) on 6 by 6 patch squares and using a B-factor of 100. The gain reference used for MotionCor2 v1.4.0 was generated by using the Sum_all_tifs program, which is distributed within the cisTEM image processing suit (80). The motion corrected micrographs were imported into cryoSPARC V3.2.0 (81), which was then used to perform patch CTF estimation and particle selection. Manually selected particles were initially extracted with a box size of 384 pixels and then used to perform 2D classification. The class averages with different views from this initial round of 2D classification were used as 2D templates for template-based particle selection in cryoSPARC, and then the selected particles were extracted and subjected to another round of 2D classification to produce improved 2D classes characterized by slightly different views. We performed iterative cycles of template-based particle selection and 2D classification until we were able to maximize the recovery of different views of particles. For template picking, we set a particle diameter of 180 Å with an overlap that did not allow any two picks to be closer than 0.4 units of particle diameter in distance. 926,176 particles were extracted with a box size of 384 pixels, following particle inspection. This stack was then subjected to downstream processing. We performed several rounds of 2D classification with this stack, leaving 466,246 particles from best 2D classes. The remaining particles were then subjected to heterogeneous refinement using an imported map of the MVV CSC intasome (EMDB-4138) (1) in cryoSPARC. The particles were split into two classes through heterogeneous refinement, and the best class (based on visual inspection and measurement of resolution) was selected to perform one round of non-uniform refinement (82). The refined map from non-uniform refinement was used as an input for the next round of heterogeneous refinement to further sort out bad particles. This iterative process, consisting of heterogeneous refinement and non-uniform refinement, was continued until we observed no further improvements to the resolution, as measured as the Fourier shell correlation (FSC) between reconstructions from pairs of independent half-sets (83) (see Fig. S10c). At this point, 147,860 particles remained and were used for the final non-uniform refinement, which was also combined with per-particle defocus refinement. The final global resolution was calculated in cryoSPARC as 3.43 Å using FSC analysis with a fixed threshold at 0.143 (Fig. S1a). The local resolution was calculated using cryoSPARC (Fig. S2a). The 3D FSC was obtained by using the 3D FSC server (https://3dfsc.salk.edu) (78) and the sampling compensation factor (SCF) and surface sampling plots were calculated using the graphical user interface tool (https://www.github.com/LyumkisLab/SamplingGui) (84, 85). Selected image analysis results are shown in Fig. S10 and summarized in Table S1.

### MVV STC cryo-EM grid preparation, data collection and image processing

A 4-μL drop of freshly prepared MVV STC at 0.15 mg/mL in 310 mM NaCl, 3 mM CaCl_2_ and 25 mM BisTris-HCl, pH 6.5 was applied onto glow-discharged lacey carbon grids coated with 3-nm carbon film (Ted Pella, product code #01824). The grids were incubated for 60 s under 100% humidity in a Vitrobot Mark IV (FEI) at 20 °C. To reduce salt concentration, the grids were blotted for 0.5 s, immediately re-hydrated with a 4-μL drop of 75 mM NaCl and 25 mM BisTris-HCl, pH 6.5 and blotted again for 3.5 s, followed by plunging into liquid ethane. Cryo-EM data were collected on a Titan Krios electron microscope (FEI) operating at 300 keV equipped with a K2 Summit direct electron detector (Gatan) using EPU software (FEI). A total of 12,679 micrograph movie stacks were acquired at a magnification of 36,232, resulting in a pixel size of 1.38 Å at the specimen level, using a total fluence of 50 e/Å^2^ spread over 50 frames with a defocus range of −1.0 to −4.5 μm.

The movies were corrected for drift and beam-induced motion applying dose weighting as implemented in MotionCor2 (79). The contrast transfer function was estimated using CTFFIND4 (86) via Relion-2.1 interface (87). Resulting frame sums were examined individually, and those containing mostly lacey carbon or showing evidence of crystalline ice contamination were discarded. A remaining 11,760 micrographs were used for further image processing (Fig. S11a). Particles, picked manually on a subset of micrographs using EMAN Boxer (88), were subjected to reference-free 2D classification. Six well-defined 2D class averages, low-pass filtered to 20 Å, were used as references for automated particle picking in Relion-2.1 of the entire dataset. Particles picked along carbon edges were removed using Eraser tool in EMAN Boxer. Remaining 2,022,321 particles were extracted, with 2-fold binning applied, and subjected to several rounds of reference-free 2D classification in Relion-2.1. A total of 684,262 particles belonging to well-resolved 2D classes (Fig. S11b) were subjected to 3D classification in Relion-2.0 into four classes without imposing symmetry (Fig. S11c); 220,096 particles belonging to the single high-resolution class were re-extracted without binning, with a box size of 320×320 pixels, and used for 3D refinement without imposing symmetry. The resulting reconstruction showed features suggesting presence of IBD bound to two symmetric positions within the CIC. To improve occupancy of LEDGF/p75, the particles were subjected to 3D classification without re-alignment with a mask focused on the IBDs and the DNA component of the STC. Prior to classification, all IN-derived signal was removed, using particle subtraction utility in Relion-2.1. The procedure allowed isolation of a 3D class harboring two IBDs and comprising 121,619 particles (Fig. S11d). Upon reverting to original non-subtracted particles, this subset was subjected to Bayesian polishing in Relion-3.1. Final reconstruction was obtained using non-uniform refinement following local and global CTF refinement (per-particle defocus and beam tilt, respectively), as implemented in cryoSPARC. To aid in model building and for illustration purposes, the map was filtered and sharpened using deepEMhancer (89) or using density modification procedure in Phenix (90). Details of EM processing statistics are given in Table S1. Resolution is reported according to the gold-standard Fourier shell correlation (FSC), using the 0.143 criterion (91, 92) (Fig. S1b), and local resolution was calculated using cryoSPARC (Fig. S2a). The 3D FSC was obtained by using the 3D FSC server (https://3dfsc.salk.edu) (78) and the sampling compensation factor (SCF) and surface sampling plots were calculated using the graphical user interface tool (https://www.github.com/LyumkisLab/SamplingGui) (84, 85).

### Real-space refinement

The STC model was constructed from the original low-resolution strand transfer complex model published previously (PDB code 5M0R) (1) and coordinates for LEDGF/p75 IBD (PDB code 3HPH) (33). The coordinates were docked into the cryo-EM map using UCSF Chimera (93). Manual adjustments were made to the model using Coot (94) and the coordinates were subjected to real-space refinement in Phenix dev-4213-000 employing *C2* NCS constraints (95). The final model has good fit to the cryo-EM map (CC=0.77) and reasonable stereochemistry as assessed using MolProbity (96). The refined STC model, with the tDNA removed, was then docked into the CSC cryo-EM map using UCSF Chimera (93). To address slight differences in the STC and CSC structures, individual domains that were not well fitted to the CSC map were docked as rigid bodies to achieve a best-fit starting model. Two (*C2*-related) NTDs (IN residues 1-35, in chains D and L) were removed from the model due to lack of supporting map. Manual adjustments were made to the model using Coot (94) and the coordinates were subjected to real-space refinement in Phenix dev-4213-000 employing *C2* NCS constraints (95). The final model has good fit to the cryo-EM map (CC=0.70) and reasonable stereochemistry as assessed using MolProbity (96). Details of the map and model statistics are given in Table S1. Model-vs-map FSCs for both structures are shown in Fig. S1c. The final cryo-EM reconstruction and fitted coordinates will be deposited with the EMDB and the PDB upon provisional acceptance of the manuscript.

### SEC-MALLS

All measurements were conducted using a Jasco chromatography system equipped with a DAWN-HELEOS II laser photometer and an OPTILAB-TrEX differential refractometer (Wyatt Technology). MVV IN was diluted to 1, 2, 4, or 8 mg/mL in 1 M NaCl, 7.5 mM 3-[(3-cholamidopropyl) dimethylammonio]-1-propanesulfonate, 2 mM DTT, 25 mM BisTris propane-HCl, pH 7.0 prior to injection of 20 µL samples onto a Superdex 200 Increase 3.2/300 column (GE Healthcare) equilibrated in 1 M NaCl, 3 mM NaN_3_, 25 mM BisTris-HCl, pH 6.5. Chromatography was performed at 20°C and a flow rate of 150 µL/min. Weight-averaged molecular masses of eluting species were calculated using the data from both detectors in ASTRA-6.1 software (Wyatt Technology).

### In vitro MVV IN strand transfer assays

Strand transfer assays were carried out as previously described (1) using double-stranded oligonucleotide mimicking the terminal 29 bp of the processed MVV U5 vDNA end (Fig. S5). The vDNA substrate was prepared by annealing synthetic HPLC-purified oligonucleotides 5’-CCGTGCAACACCGGAGCGGATCTCGCA and 5’-GCTGCGAGATCCGCTCCGGTGTTGCACGG. Reactions were initiated by addition of 1.1 μM MVV IN to 1.5 μM LEDGF/p75, 0.75 μM vDNA substrate and 7.5 ng/μL supercoiled target pGEM9zf DNA (Promega) in 25 mM BisTris-HCl, pH 6.0, 40 mM KCl, 5 mM MgSO_4_, 5 μM ZnCl_2_ and 2.5 mM DTT, in a final volume of 40 μL. Reactions were allowed to proceed for 1 h at 37°C and were stopped by addition of 25 mM EDTA and 0.5% SDS. DNA products, deproteinized by digestion with proteinase K (Thermo Fisher Scientific) and ethanol precipitation, were dissolved in 20 μL of deionized water. Products were analyzed by electrophoresis through 1.5% agarose gels in Tris-Acetate-EDTA buffer and detected by staining with ethidium bromide. Strand transfer products were quantified using real-time PCR on the deproteinized DNA samples using primers 5’-CCGGCTTTCCCCGTCAAGCT and 5’-ACACCGGAGCGGATCTCG, annealing within pGEM9zf plasmid and vDNA, respectively. The real-time quantitative PCR reactions were carried out in triplicates, with 5 μM of each primer, 1 μL strand transfer product DNA (diluted 1:2,500) in PowerUp SYBR-Green master mix (Thermo Fisher Scientific) and a total reaction volume of 20 μL. Relative quantities of the strand transfer products were estimated using standard curves, generated by serial dilutions of an upscaled sample with WT IN.

### TIRF microscopy and photobleaching

All experiments were performed at room temperature in a microfluidic flow cell, functionalized with partially biotinylated polyethylene glycol (mixture of mPEG-SVA-5000 and Biotin-PEG-SVA-5000 at 50:1 mass ratio; Lysan) and assembled as previously described (97). Prior to experiments, the biotinylated surface was coated for 10 min with 0.2 mg/mL *Streptomyces avidinii* streptavidin (Sigma-Aldrich) in PBS. All buffers and solutions were thoroughly degassed immediately before use. Flow was controlled by an automated syringe pump (Pump 11 Elite; Harvard Apparatus).

Flow cells were mounted on a Nikon Eclipse Ti inverted microscope, equipped with a 100x high numerical aperture TIRF objective (SR Apo TIRF 100x 1.49 Oil; Nikon). Cy3 and Surf649 dyes were excited with 561 and 640 nm lasers (LU-N4 laser unit; Nikon), respectively, at 10% of maximum power. Emitted fluorescence was recorded using a 512 x 512 pixel, back illuminated, electron-multiplying charge-coupled-device camera (iXon DU-987, Andor Technology; 3 MHz pixel readout rate, 14-bit digitization and 300x electron multiplier gain) and a frame rate of 5 Hz. The pixel size was 160 x 160 nm.

CSC intasomes (125 μL of 4 pM solution) containing biotinylated and Cy3-labelled vDNA (prepared by annealing 5’-Cy3-CCGTGCAACACCGGAGCGGATCTCGCA and 5’-GCTGCGAGATCCGCTCCGGTGTTGCACGG-TEG-Biotin) in Buffer A (1 M NaCl, 3 mM CaCl_2_, 1 mg/mL BSA, 1 mg/mL casein, and 25 mM BisTris-HCl, pH 6.5) was drawn into the streptavidin-coated flow cell pre-equilibrated with Buffer A at a 25 μL/min flow rate and incubated for further 10 min. Excess intasome was washed off with 250 μL Buffer A at a flow rate of 50 μL/min. Next, 125 μL of a 33-nM solution of LEDGF/p75-Surf649 in Buffer B (0.5 M NaCl, 3 mM CaCl_2_, 1 mg/mL BSA, 1 mg/mL casein, and 25 mM BisTris-HCl, pH 6.5) or C (0.2 M NaCl, 3 mM CaCl_2_, 1 mg/mL BSA, 1 mg/mL casein, and 25 mM BisTris-HCl, pH 6.5) were introduced into the flow cell (pre-equilibrated with the same buffer) at a flow rate of 25 μL/min. After a 10-min incubation, excess LEDGF/p75-Surf649 was washed off with 250 μL of the same buffer. At this point, the following imaging sequence was implemented for three individual fields of view (512 x 512 pixels) per buffer condition: Cy3, for 5 frames, Surf649 for 3 min of continuous sampling (resulting in over 95% photobleaching within the range of the TIRF field), followed by Cy3 for 5 frames.

All data sets were initially processed with NIS Elements (Nikon) for noise reduction (“advanced denoising” at a value of 5 for both analyzed channels) and background subtraction (“rolling ball” algorithm with r = 0.48 μm). For each field of view, Cy3 (intasome) spots were detected in the 5-frame initial dataset, using “bright spot detection” (0.78 μm radius and 14.5 contrast) within NIS Elements. On average, 400 spots were detected per 512 x 512 pixel field of view. Next, for each identified intasome spot, a Surf649 (LEDGF/p75) fluorescence trace was obtained from the 3-min continuous sampling acquisition. Spots with Cy3 fluorescence intensity above 200 arbitrary units were not considered for analysis as they are likely to represent overlapping intasome complexes (the 200 arbitrary units threshold was chosen based on Cy3 intensity distribution for all analyzed data). Similarly, data with no Surf649 photobleaching steps or unstable bleaching profiles were excluded from further analysis. In order to obtain the number of Surf649 photobleaching steps per intasome, firstly, pairwise intensity differences were calculated for each photobleach trace: *ΔI_ij_ = ΔI(t_i_) − ΔI(t_j_)* for all data pairs, for which *t_i_ > t_j_*, followed by local maxima identification. These calculations were performed in MATLAB (MathWorks). For each salt condition, the number of photobleaching steps per intasome was statistically analyzed for all three fields of view. Prism 7 (GraphPad) was used for statistical analysis and data plotting.

### Cell lines and tissue culture

Cells were cultured at 37°C in 5% CO_2_ atmosphere in Dulbecco’s modified Eagle medium (DMEM, Life Technologies) supplemented with 10% heat-inactivated fetal bovine serum (FBS) and antibiotic/antimycotic solution (Sigma-Aldrich). LKO is a LEDGF-null cell line, generated via TALEN-mediated *PSIP1* gene disruption in HEK293T cells (51). Cells additionally null for HRP2 (LHKO) were generated from LKO cells using CRISPR-Cas9 guide RNAs targeting exon 2 (5’-ACCCAACAAGTACCCCATCTTTTTC) and exon 15 (5’-CGACCGGCAGGAGCGCGAGAGGG), resulting in deletion of most of *HDGFL2* (26 kb). Gene disruption was verified by genomic DNA sequencing, and the absence of detectable HRP2 protein was verified by immunoblotting. For ectopic expression of ovine LEDGF/p75, LHKO cells were transduced with murine leukemia virus virus-based retroviral vector pQ-*Oa*LEDGF-IRES-PuroR and selected in the presence of 1 μg/mL puromycin. To construct this vector, a DNA fragment encoding unaltered full-length ovine LEDGF/p75 (GenBank accession code FJ497048), PCR-amplified from a sheep peripheral blood mononuclear cell cDNA library (33) using primers 5’-GCGCATGCGGCCGCAGACACCATGACTCGCGACTTCAAACCTGGGGACC and 5’-GGCGGGATCCCTAGTTATCTAGTGTAGAATCCTTCAGAG, was ligated between *Not*I and *Bam*HI restriction sites of pQCXIP (Clontech).

CPT3 was a subclone of CPT-Tert, an ovine choroid plexus cell line immortalized by co-expression of simian virus 40 large T antigen and human telomerase RT (52). For genome modification of CPT3 cells, guide RNAs were prepared using CRISPR RNA targeting sheep *PSIP1* or *HDGFL2* genes (Table S3) and Alt-R CRISPR-Cas9 tracrRNA (IDT) each at a final concentration of 50 μM. The ribonucleoprotein complexes were prepared by incubating 30 μM guide RNA and 24.8 μM recombinant Alt-R Cas9 Nuclease V3 (IDT) for 20 min at room temperature. CPT3 cells (1.5×10^6^) were resuspended in 70 μl SE Cell Line Nucleofector solution (Lonza) and 18-36 μl ribonucleoprotein complexes were added along with 4 μM Alt-R Cas9 electroporation enhancer (IDT) to a nucleofection chamber (Lonza). Cells were electroporated using the nucleofector program EN-138 and gently resuspended in DMEM, before plating in a 6-well dish. The knockout cell lines were generated by transfection of CPT3 cells with ribonucleoprotein complexes containing crRNAs 2, 3, and 4 (LKO1 and LKO2); crRNAs 2 and 3 (LKO3); crRNAs 2 and 4 (LKO4); 2, 3, 5, 6, 7, and 8 (LHKO1); and 2, 4, 5, 6, 7, and 8 (LHKO2). Target sequences for CRISPR RNA are given in Table S5.

### Western blotting

The following primary antibodies were used: rabbit polyclonal anti-LEDGF/p75 (Bethyl Laboratories; product code A300-848A), rabbit polyclonal anti-HRP2 (Novus Bio; product code NBP2-47438), affinity-purified polyclonal rabbit anti-MVV capsid/p24 antibody (Thermo Fisher Scientific; custom product code HAB2110A), and horseradish peroxidase-conjugated mouse monoclonal anti-β-actin antibody (Cell Signaling Technology, product code 5125). Blots probed with anti-LEDGF/p75, anti-HRP2, and anti-capsid/p24 antibodies were developed following incubation with horseradish peroxidase-conjugated goat anti-rabbit IgG antibody (BioRad; product code 1706515) or IRDye 800CW conjugated goat anti-rabbit IgG antibody (LI-COR; product code 925-32211) for detection by chemiluminescence, using Imager 600 RGB (GE Healthcare) following incubation with ECL prime reagent (GE Healthcare) or by fluorescence, using Odyssey imager (LI-COR), respectively.

### MVV vectors

The packaging construct pCAG-MV-GagPol-CTEx2 (44, 45) is based on MVV isolate EV1 (98). To enable mutagenesis, the construct was modified by flanking its IN-coding region with *Age*I and *Xho*I restriction sites. To this end, a DNA fragment spanning *Pas*I and *Dra*III sites of pCAG-MV-GagPol-CTEx2 was PCR-amplified and ligated between *Bam*HI and *Xho*I sites of pBluescript II KS(+) (Stratagene). *Age*I and *Xho*I restriction sites flanking the IN-coding region were introduced by silent mutagenesis, and the modified DNA fragment was ligated between *Pas*I and *Dra*III sites of the packaging construct. Next, a DNA fragment encoding IN from KV1772 MVV isolate (74) was ligated between *Age*I and *Xho*I sites of the modified packaging construct, resulting in pCAG-MV-GagPol-IN^KV1772^-CTEx2. Importantly, these alterations did not affect infectivity of the MVV vector particles (Fig. S6c) and streamlined the genetic analyses due to availability of mutant KV1772 IN expression constructs. EV1 and KV1772 INs share 86.2% amino acid sequence identity and 92% similarity. Variants of the modified packaging construct were obtained by replacing the WT IN coding region with the corresponding mutant sequences. MVV transfer vector pCVW-CG-Luc, which encodes for firefly luciferase, was constructed by overlapping PCR. Initial PCR amplicons carrying the CMV promoter and firefly luciferase were amplified from MVV transfer vector pCVW-CG and pHI.Luc (99), respectively. A single linked fragment, which was produced by a second round of PCR, was digested *Blp*I and *Bgl*II, and then ligated with *Blp*I/*Bgl*II-digested pCVW-CG.

MVV vector particles were produced by transfection of HEK293T cells with four-component MVV vector system (44, 45) using polyethyleneimine (PEI). Briefly, HEK293T cells were plated in 15-cm Petri dishes to achieve ∼80% confluency on the day of transfection. The cell medium was then replaced with 15 mL OptiMEM reduced serum medium (Life Technologies). PEI-DNA complexes were prepared by combining 58.5 μL 0.1% PEI (w/v in PBS, Sigma-Aldrich product number #408727) with the four-plasmid mixture containing 10.5 μg pCAG-MV-GagPol-IN^KV1772^-CTEx2 (WT or mutant MVV packaging construct), 15.75 μg pCVW-CG-Luc (MVV transfer vector encoding firefly luciferase gene reporter) or pCVW-CG, which encodes for GFP, 3.5 μg pCMV-VMV-Rev (MVV Rev expression construct), and 5.25 μg pMD2.G (VSV-G expression construct, a gift from D. Trono) pre-diluted into 2.5 mL OptiMEM. Following incubation at room temperature for 15-20 min, PEI-DNA complexes were added dropwise to cells. Next day, medium was replaced with fresh DMEM supplemented with 10% FBS. Cell culture supernatant containing viral particles was harvested 36-48 h post-transfection and pre-cleared by filtration through a 0.45-μm filter. Viral particles were pelleted at 30,000 rpm for 1.5 h at 25°C in an Optima XPN ultracentrifuge using an SW 32 Ti rotor (Beckman Coulter), resuspended in 500 μL DMEM supplemented with 10% FBS, and stored in small aliquots at −80°C. Prior to infections, viruses were treated with 0.12 U/μL TURBO DNase (Thermo Fisher Scientific) at 37 °C for 1 h. Infections of HEK293T, CPT3, and derivative cell lines were carried out in 48-well plates with the virus quantity corresponded to 0.5 mU of associated RT activity; 2 d post-infection, cells were expanded into 6-well plates, and passaged (1:2 - 1:10) 5 d post-infection. Cells were harvested 7 d post-infection.

### Quantification of vector particles using product-enhanced RT assay

Virus-associated RT activity was measured using real-time quantitative PCR using a published method (100) and adapted for the TaqMan platform (17). Viral particles were disrupted by combining with equal volume of lysis buffer (typically 5 μl) containing 0.8 U/μL RiboLock RNase Inhibitor (Thermo Fisher Scientific) in 50 mM KCl, 40% (v/v) glycerol, 0.25% (v/v) Triton-X100, 100 mM Tris-HCl, pH 7.4. Following incubation at room temperature for 10 min, viral lysates were diluted with 9 volumes of core buffer (20 mM KCl, 5 mM ammonium sulfate, and 20 mM Tris-HCl, pH 8.3). Each real-time PCR reaction mixture comprised 12.5 μL TaqMan Gene Expression Master Mix (Thermo Fisher Scientific), primers 5’-TCCTGCTCAACTTCCTGTCGAG and 5’-CACAGGTCAAACCTCCTAGGAATG (each at the final concentration of 0.5 μM), 0.2 μM TaqMan probe 5’-[6FAM]-CGAGACGCTACCATGGCTATCGCTGTAG-[TAMRA], 5 U RiboLock RNase inhibitor, 100 ng phage MS2 phage RNA (Sigma-Aldrich), and 2 μL diluted viral lysate in a final volume of 25 μL. Reactions were assembled in MicroAmp Optical 96-well reaction plates and carried out in a QuantStudio-7 Flex real-time PCR instrument (Applied Biosystems). PCR conditions were as follows: 42°C for 20 min (reverse transcription step), 50°C for 2 min, 95°C for 10 min (activation of the host start Taq DNA polymerase), followed by 40 cycles of PCR (denaturation at 95°C for 15 sec and extension at 60°C for 1 min). All PCR reactions were performed in triplicate. Standard curves were generated using recombinant HIV-1 RT (Merck Millipore). The relative viral quantities were calculated based on the standard curve generated using QuantStudio-7 systems software (Applied Biosystems).

### Luciferase assay and flow cytometry

To measure luciferase activity, cells were washed in phosphate-buffered saline and lysed in 10 mM DTT, 10 mM EDTA, 50% glycerol, 5% triton X-100, and 125 mM Tris-HCl, pH 7.4. Following a single freeze-thaw cycle, insoluble proteins were pelleted by centrifugation at 21,000 *g*. Total protein content of each pre-cleared sample was determined using Pierce bicinchoninic acid assay protein assay (Thermo Fisher Scientific). To measure luciferase activity, 10 μL pre-cleared cell lysate was combined with 200 μL assay reagent containing 1 mM D-luciferin (BD Biosciences, product #556878), 3 mM adenosine triphosphate, 15 mM MgSO_4_, and 30 mM sodium 4-(2-hydroxyethyl)-1-piperazineethanesulfonate, pH 7.8. Luminescence measured using a EnVision 2102 plate reader (Perkin Elmer), was normalized by the total protein content in each precleared lysate.

Flow cytometry was used to count cells infected with GFP reporter viruses. Cells were harvested by trypsinization, fixed in 6.5% (w/v) formaldehyde in PBS and analyzed for GFP expression using a Fortessa flow cytometer (BD Biosciences). GFP was excited at 495 nm and emission was detected with a 530/30 band pass filter. Data were acquired using BD Diva software (BD Biosciences) and analyzed using FlowJo v13. Live cells (population P1) were initially gated using the area plot of forward scatter (FSC-A) versus side scatter (SSC-A), to separate them from cell debris. Singe cells (population P2) were then differentiated from doublets in population P1 by gating on FSC-A vs forward scatter height (FSC-H). 10,000 single cells were analyzed for GFP expression by gating for 530/30 blue and FSC-H.

### Quantification of vDNA synthesis in infected cells

Total DNA from infected cells was extracted and purified using DNeasy Blood and Tissue kit (Qiagen) or Quick-DNA Microprep Kit (Zymo Research) at specified times post-infection with WT or mutant MVV vectors. MVV vDNA was quantified using real-time PCR with primers 5’-GATGGTTAAGTCATAACCGCAGATG and 5’-GGTGTCTCTCTTACCTTACTTCAGG, designed to amplify a fragment within the 5’ region of the vDNA, spanning the U3-*gag* junction. Because the transfer vector construct pCVW-CG-Luc lacks a complete 5’ LTR (expression in transfected cells is driven by the CMV promoter), it cannot template amplification of the PCR product. Therefore, the strategy allowed the selective quantification of late reverse transcription products even in the presence of background plasmid DNA that may have persisted despite DNase treatment of vector particles prior to infection. Amplification was detected using TaqMan probe 5’-[FAM]CGTGGGGCTCGAC AAAGAATC[TAMRA]. To generate standard curves, the PCR product inserted into pCR4-TOPO vector (Thermo Fisher Scientific) was employed as template. Each real-time PCR reaction mixture comprised 10 μL TaqMan Gene Expression Master Mix (supplied as a 2x concentrate; Thermo Fisher Scientific), primers (each at the final concentration of 0.1 μM), 0.2 μM TaqMan probe, and 4 μL DNA sample (corresponding to 100 ng total DNA) in a final volume of 20 μL. Reactions were assembled in MicroAmp Optical 96-well reaction plates and carried out in a QuantStudio 7 Flex real-time PCR instrument (Applied Biosystems). PCR conditions were as follows: 50°C for 2 min, 95°C for 10 min (activation of the hot start Taq DNA polymerase), followed by 40 cycles of PCR (95°C for 15 sec, 60°C for 1 min, 72°C for 30 sec). All PCR reactions were performed in triplicate.

### Mapping MVV vector integration sites in human and ovine cells

Genomic DNA isolated from cells 5 d post-infection with MVV vectors was processed for linker-mediated PCR, and the amplified viral LTR-chromosome junctions were sequenced as previously described (1, 101). Briefly, genomic DNA, digested with *Mse*I overnight at 37°C, was ligated to a double-stranded DNA linker containing 5’-TA overhang overnight at 12°C. Customized DNA linkers were used in conjunction with barcoded primers to aid in multiplexing and prevent crosstalk between samples. The first round and the nested MVV U5 primers were 5’-CTAATTCCGTGCAACACCG and 5’-AAT GAT ACG GCG ACC ACC GAG ATC TAC ACT CTT TCC CTA CAC GAC GCT CTT CCG ATC T<u>NN NNN N</u>CAACA CCG GAG CGG ATC, respectively (underlined sequence represents the 6 nucleotides barcode specific to LTR primer for multiplexing). The nested PCR primers contained Illumina adaptor sequences appended at the 5’ ends. The PCR products, multiplexed on a single lane of a flow cell, were subjected to 150-bp paired-end sequencing on a HiSeq-4000 Illumina instrument (GENEWIZ, Boston, MA, USA).

Paired-end Illumina sequence reads, cropped to remove linker- and viral U5 DNA-derived sequences, were aligned with the hg38 version of human or the oviAri4 version of sheep genome using BWA MEM (102). The results were parsed using SAMtools (103) to extract unique, high-confidence alignments (samtools view -F 4 -F 256 -q 1). Only reads matching the host genome sequence immediately downstream of the processed vDNA U5 end were considered for further analysis. Genomic coordinates of the unique integration sites were converted to the browser extensible data format (with each interval corresponding to the middle dinucleotide of an integration site). Distributions of the integration sites with respect to various genomic features (http://genome.ucsc.edu/cgi-bin/hgTables), cLADs (104), SPADs (55) as well as local gene density and nucleotide content at the sites of integration were calculated using BEDtools (105). Expression data averaged across sheep brain tissues (106) was used to approximate gene activity in CPT-Tert cells; HEK293T gene expression data were from Ref. (107). Local gene density and nucleotide content comparisons were computed using Wilcoxon rank sum test, and significance of all other comparisons were calculated with Fisher’s exact test. Sequence logos were generated using WebLogo (108). HIV-1 integration sites in HEK293T and LKO cells were from published datasets (56, 109).

## Acknowledgements

We thank Massimo Palmarini for CPT-Tert cells, Didier Trono for a generous gift of pMD2.G, Ron Vale for His_10_-PS-SNAPf vector, Goedele Maertens for sharing the luciferase assay protocol and critical reading of the manuscript, Massimo Pizzato for advice on RT assays and nucleofection, and P. Walker and A. Purkiss for computer and software support. This work was funded by US National Institutes of Health grants P50 AI150481 (PC and ANE), R01 AI070042 (ANE), and U54 AI150472 (DL); US National Science Foundation CAREER MCB-2048095 and the Margaret T. Morris Foundation (DL); the Spanish Ministry of Science and Innovation PID2019-108850RA-I00 (JV); and the Francis Crick Institute (PC, HY, and IAT), which receives its core funding from Cancer Research UK (FC001061, FC001221, FC001178), the UK Medical Research Council (FC001061, FC001221, FC001178), and the Wellcome Trust (FC001061, FC001221, FC001178).

## Author contributions

ABC optimized production of MVV STC intasomes and vitrified samples for cryo-EM; ABC and PC refined the STC cryo-EM structure; VC, ABC, ZS, GJB, and NC, produced recombinant proteins; VC, conducted *in vitro* activity assays, studied phenotypes of the MVV vector mutants, generated CPT3-LKO and LHKO cell lines, and prepared samples for MVV integration site sequencing; VC and IAT conducted SEC-MALLS experiments; DTG and HY did TIRF microscopy and photobleaching analyses; ZS and DL prepared CSC intasomes, collected cryo-EM data and refined the structure; PKS, ANE, and PC mapped and analyzed integration site distributions; AN collected STC cryo-EM data; PC and JV post-processed cryo-EM maps; VEP assembled and refined the CSC and STS models; RKM and DJG constructed single-cycle MVV virus system; WL constructed luciferase reported vector; HJF and EMP established LKO and LHKO cell lines.

---

**Supplementary Figure S1.**
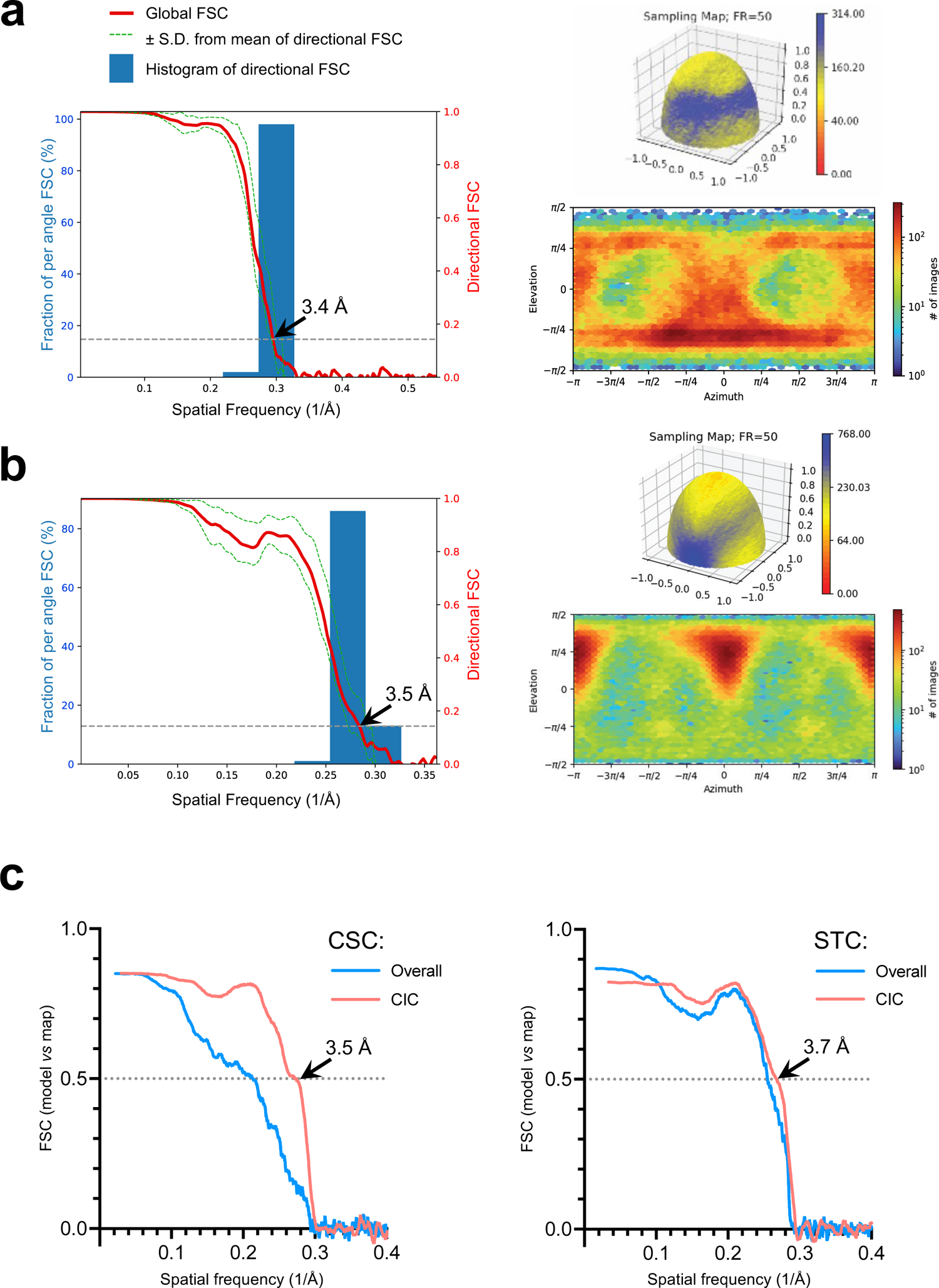
Directional resolution of cryo-EM reconstructions, particle views bias, and model fit indicators. **(A)** Directional resolution metrics and estimated projection distributions for the MVV CSC. Left: the output generated by the 3DFSC software (78) showing the global FSC curve (thick red line), boundaries of the directional FSCs (± 1 standard deviation, dotted green lines), and a histogram of directional FSC values (blue bars). Estimated nominal resolution using the fixed FSC 0.143 threshold is indicated with a black arrowhead. Top right and bottom right show the surface sampling plot (85) and estimated Euler angle distribution for the set of particles contributing to the 3D reconstruction. **(B)** Same as in panel A for MVV STC. **(C)** Model vs map FSC plots for MVV CSC (left) and STC (right). Blue and orange lines represent FSCs for the entire model and the CIC, respectively.

**Supplementary Figure S2.**
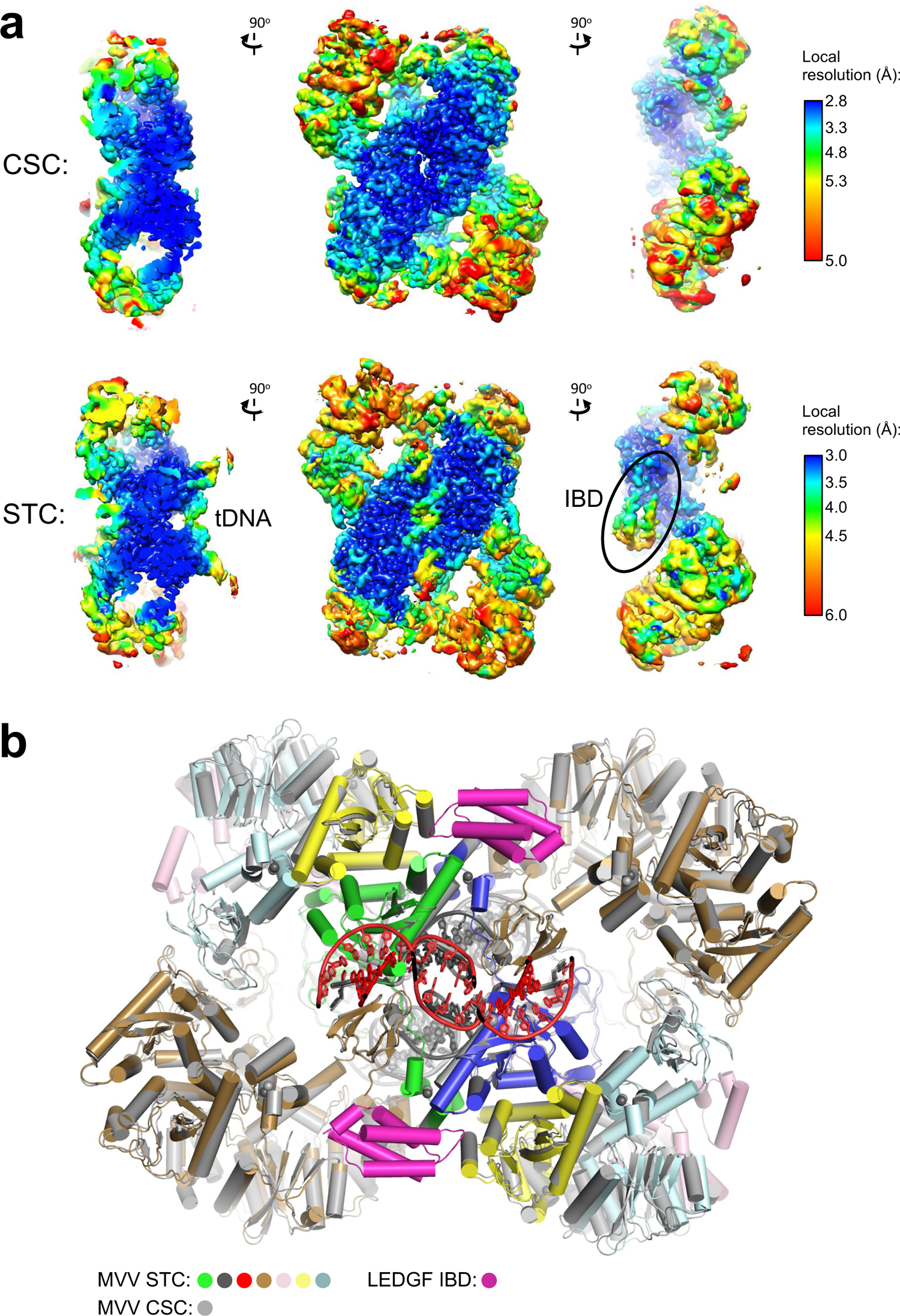
Local resolutions the cryo-EM reconstructions and comparison of the final CSC and STC models. **(A)** Local resolution distributions for CSC (top) and STC (bottom) cryo-EM reconstructions indicated with color code corresponding to the key on the right. The maps are shown in three orientations; volumes on the left are shown as orthoslices. The map features corresponding to LEDGF/p75 IBD are indicated with a black oval. **(B)** Superposition of the refined CSC and STC models. The protein subunits are shown as cartoons, with helices as tubes and DNA as sticks. STC is colored as in Fig. 1A and CSC in grey.

**Supplementary Figure S3.**
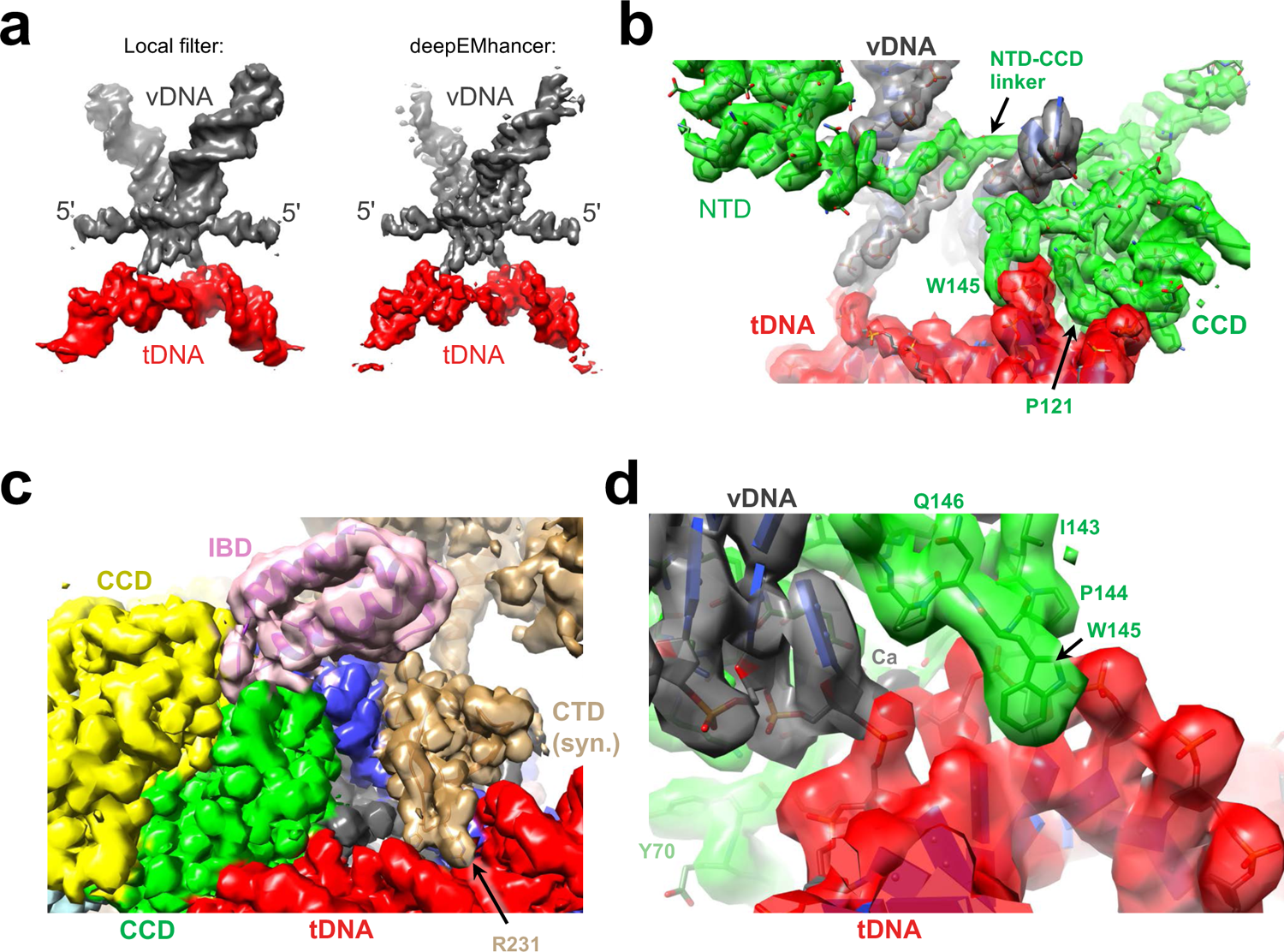
Examples of the STC cryo-EM map. **(A)** The region corresponding to the vDNA-tDNA synapse from the STC cryo-EM map sharpened using local filtering in cryoSPARC (left) or using DeepEMhancer (89) (right). 5’-unpaired vDNA nucleotides are indicated to the left. **(B-D)** Selected regions of the STC volume sharpened using DeepEMhancer. Selected structural elements, including the NTD-CCD linker (panel B), and amino acid residues are indicated.

**Supplementary Figure S4.**
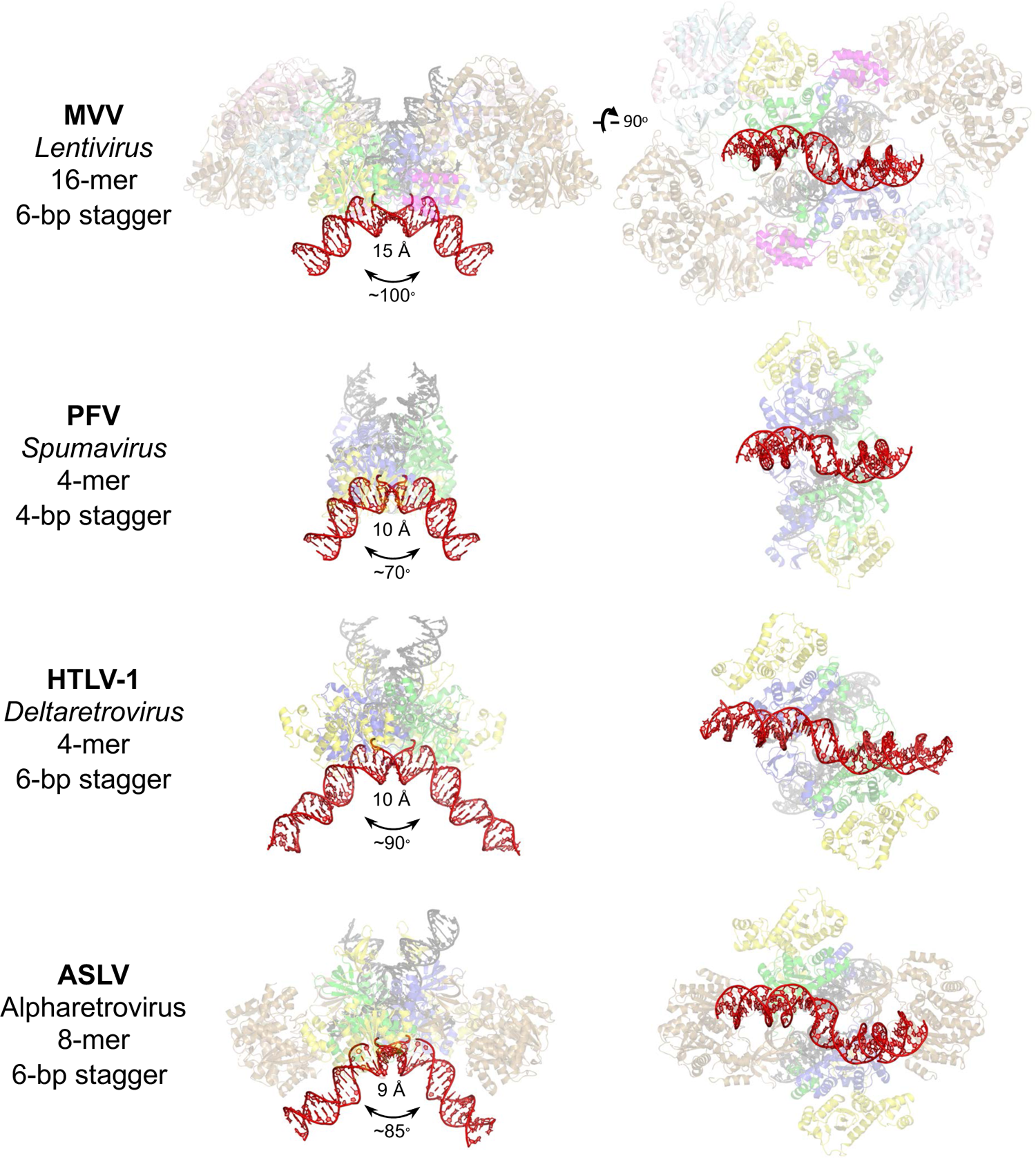
Conformations assumed by tDNA within diverse retroviral STCs. Protein subunits and DNA chains in each structure are shown in cartoons and sticks, respectively. The tDNA portions are shown opaque (red), with the rest of the structures semi-transparent. The STC structures shown are from MVV (this work), PFV (PDB ID 4BAC, Ref. (10)), HTLV-1 (PDB ID 6VOY, Ref. (15)), and ASLV (PDB ID 5EJK, Ref. (12)). Each structure is shown in two orthogonal orientations. The species, genus, corresponding IN multimeric state, and integration stagger (bp) are reported to the left. The minor groove width at the site of integration and the angle between tDNA arms are reported under each left panel.

**Supplementary Figure S5.**
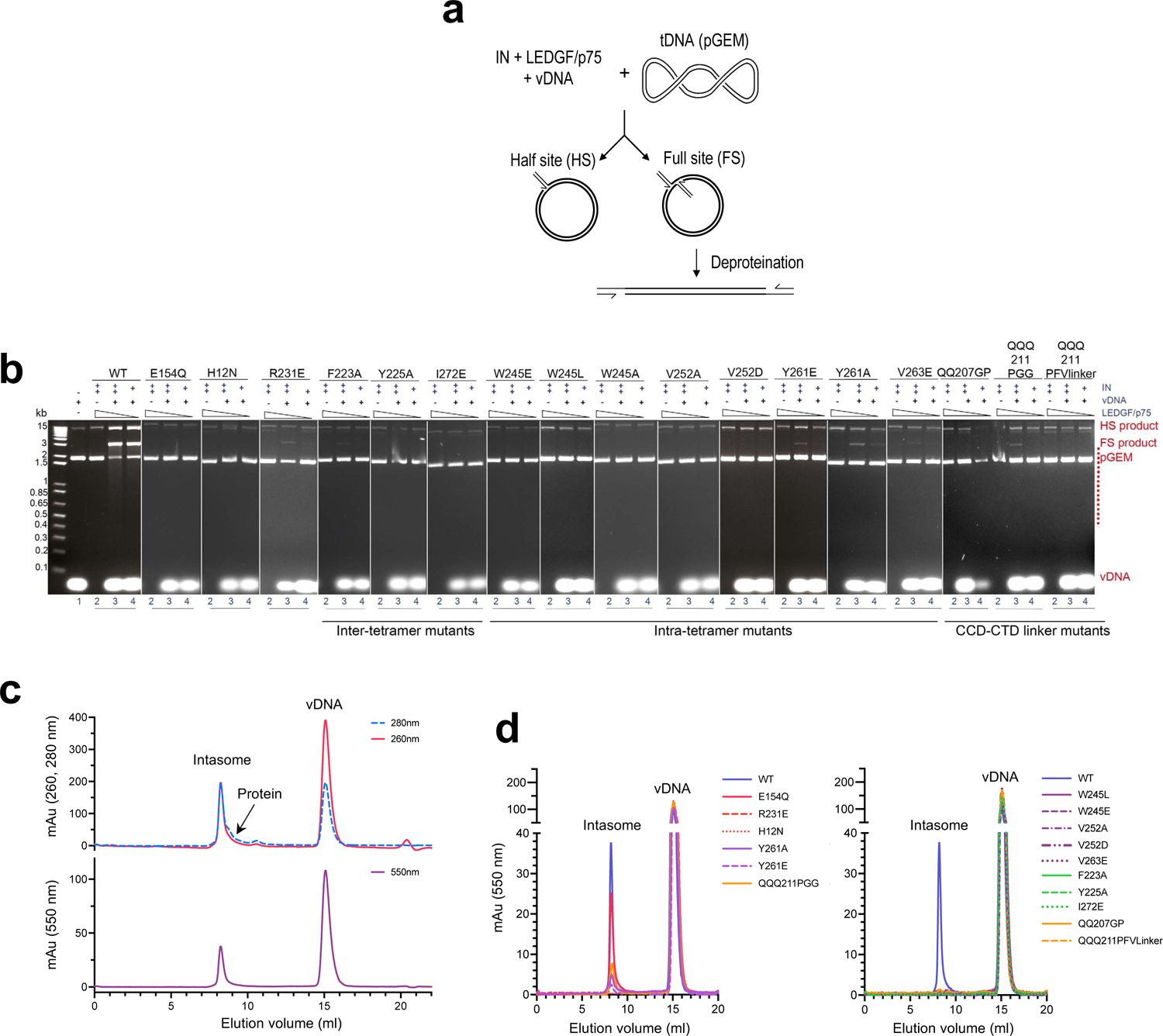
*In vitro* activities of MVV IN variants. **(A)** Schematic of the strand transfer assay, which utilizes double-stranded oligonucleotides matching the processed 3’ end (U5) of the MVV RT and supercoiled plasmid (pGEM) as mimics of vDNA ends and target DNA, respectively. The assay allows detection of two types of strand transfer products: full-site, resulting from insertion of pairs of vDNA ends into opposing strands of tDNA and half-site, resulting from insertion of a single vDNA end. When resolved by agarose gel electrophoresis, the full- and half-site products co-migrate with linearized and open-circular forms of the plasmid, respectively. Note that multiple full-site strand transfer events result in fragmentation of target DNA, giving rise to smearing that becomes pronounced at higher MVV IN and LEDGF/p75 inputs. **(B)** Strand transfer activities of MVV IN variants. WT and mutant INs at a concentration of 1.1 µM (lanes 2 and 3) or 0.55 µM (lanes 4) were incubated with 1.5 µM (lanes 2 and 3) or 0.75 µM (lanes 4) LEDGF/p75, with (lanes 3 and 4) or without (lanes 2) 20 µM vDNA in the presence of 7.5 ng/µL pGEM target DNA. Lane 1 contained a mock reaction without IN and LEDGF/p75. Deproteinized reaction products were separated in 1.5% agarose gels and detected by staining with ethidium bromide. Migration positions of the reaction products, vDNA and pGEM are indicated on the right of the gel; positions of DNA molecular size markers (kb) are shown on the left. **(C)** Analysis of CSC intasome assembly containing WT MVV IN and Cy3-labeled vDNA oligonucleotide by size exclusion chromatography. UV (wavelengths 260 and 280 nm) and visible light (550 nm) absorbance of the eluate is plotted on the top and bottom, respectively. Elution positions of the intasome, protein (IN-LEDGF complexes) and vDNA are indicated. **(D)** Elution profiles of intasome assembly reactions containing WT and two sets of mutant MVV INs and Cy3-labeled vDNA. Chromatograms on the left represent full elution profiles of the assembly reactions shown in Fig. 3D.

**Supplementary Figure S6.**
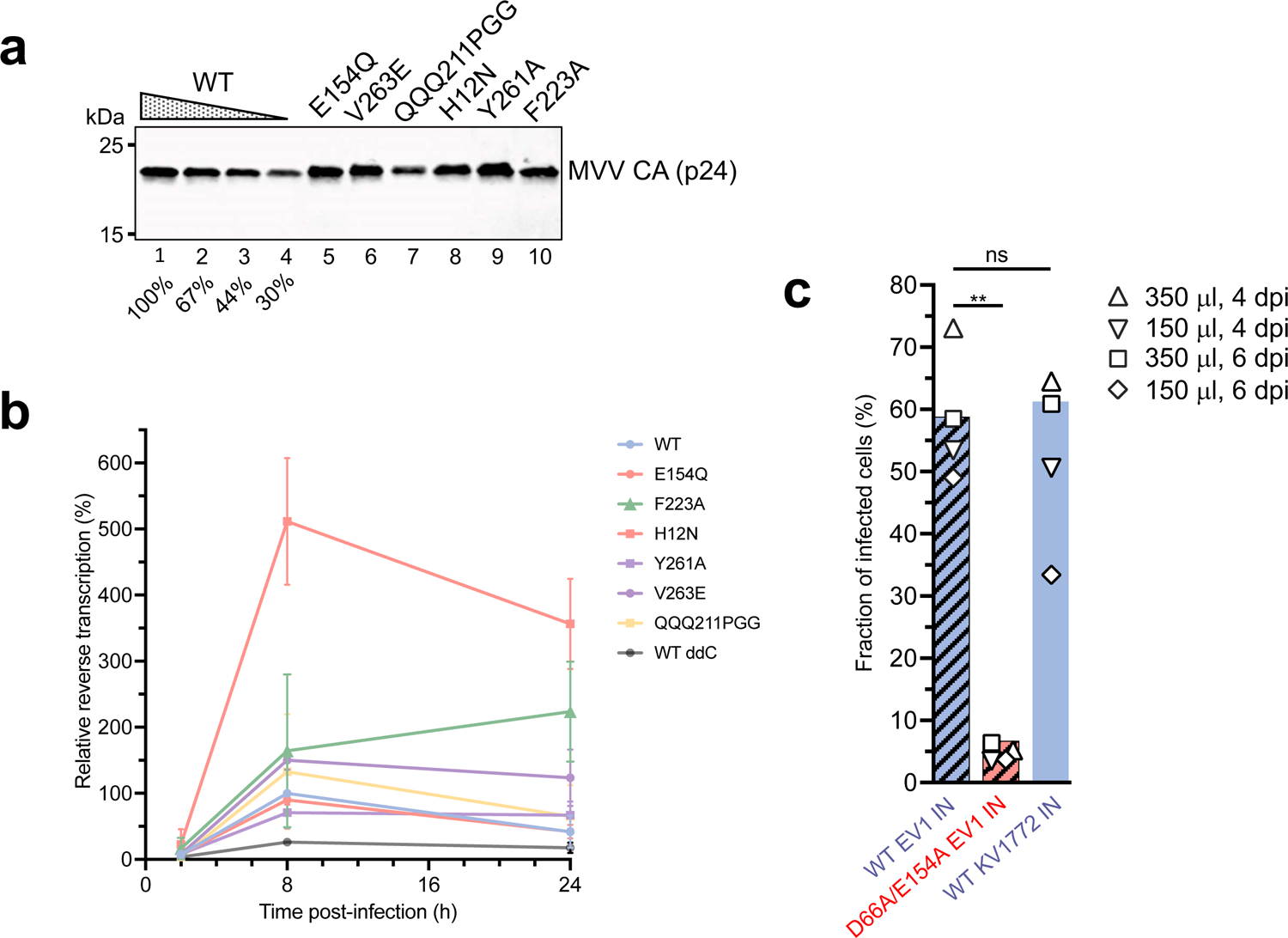
**(A)** MVV vector preparations containing 0.5 mU associated RT activity were separated in lanes 1 (WT), and 5-10 (for indicated mutants). Lanes 2, 3, and 4 contained serial 1.5-fold dilutions of the WT sample. Mature MVV capsid/p24 protein was detected by Western blotting with a rabbit polyclonal antibody and IRDye 800CW-conjugated secondary antibody. **(B)** Levels of late reverse transcription products measured in HEK293T cells at 2, 8 and 24 h post-infection with WT and IN mutant MVV vectors. Line plots represent means, and standard deviations were derived from three biological replicates. The grey line reports vDNA levels in cells infected with WT MVV in the presence of 100 µM ddC. **(C)** Infectivity of GFP-reported MVV vectors produced using Gag-Pol constructs with the IN-coding region derived from EV1 or KV1772 MVV isolate. HEK293T cells were infected with equal 350 µL (corresponding to 16.8 mU of associated RT activity) or 150 µL of WT or active site mutant (D66A/E154A) EV1, or WT KV1772 IN vector. GFP-positive cells were counted 4 or 6 d post-infection by flow cytometry. Results of four individual measurements are indicated with symbols, and bars represent fractions of GFP-expressing cells 4 days post-infection with 150 µL virus. Statistical significance was tested using paired two-tailed Student’s t-test (ns, non-significant; **, *p*<0.01).

**Supplementary Figure S7.**
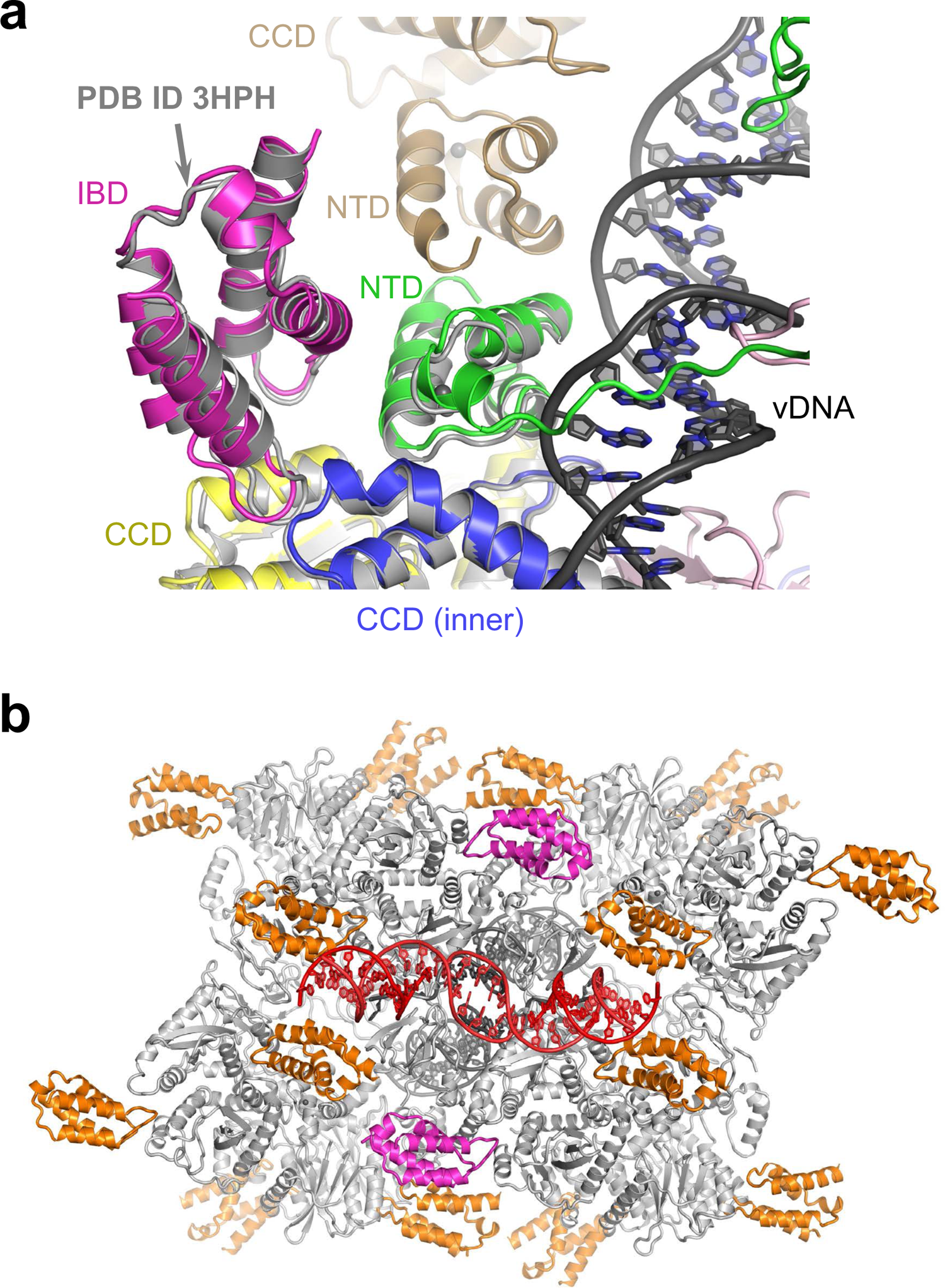
MVV intasome - LEDGF/p75 IBD interactions. **(A)** Superposition of the MVV STC (shown as cartoons and colored as in Fig. 1A) and MVV IN-LEDGF/p75 co-crystal structure (grey cartoons; PDB ID 3HPH) (33). **(B)** A model of MVV intasome (grey) with every potential LEDGF/p75 IBD binding site occupied. The two IBDs found in the STC structure are shown in magenta, while the fourteen IBDs absent in the STC reconstruction in orange.

**Supplementary Figure S8.**
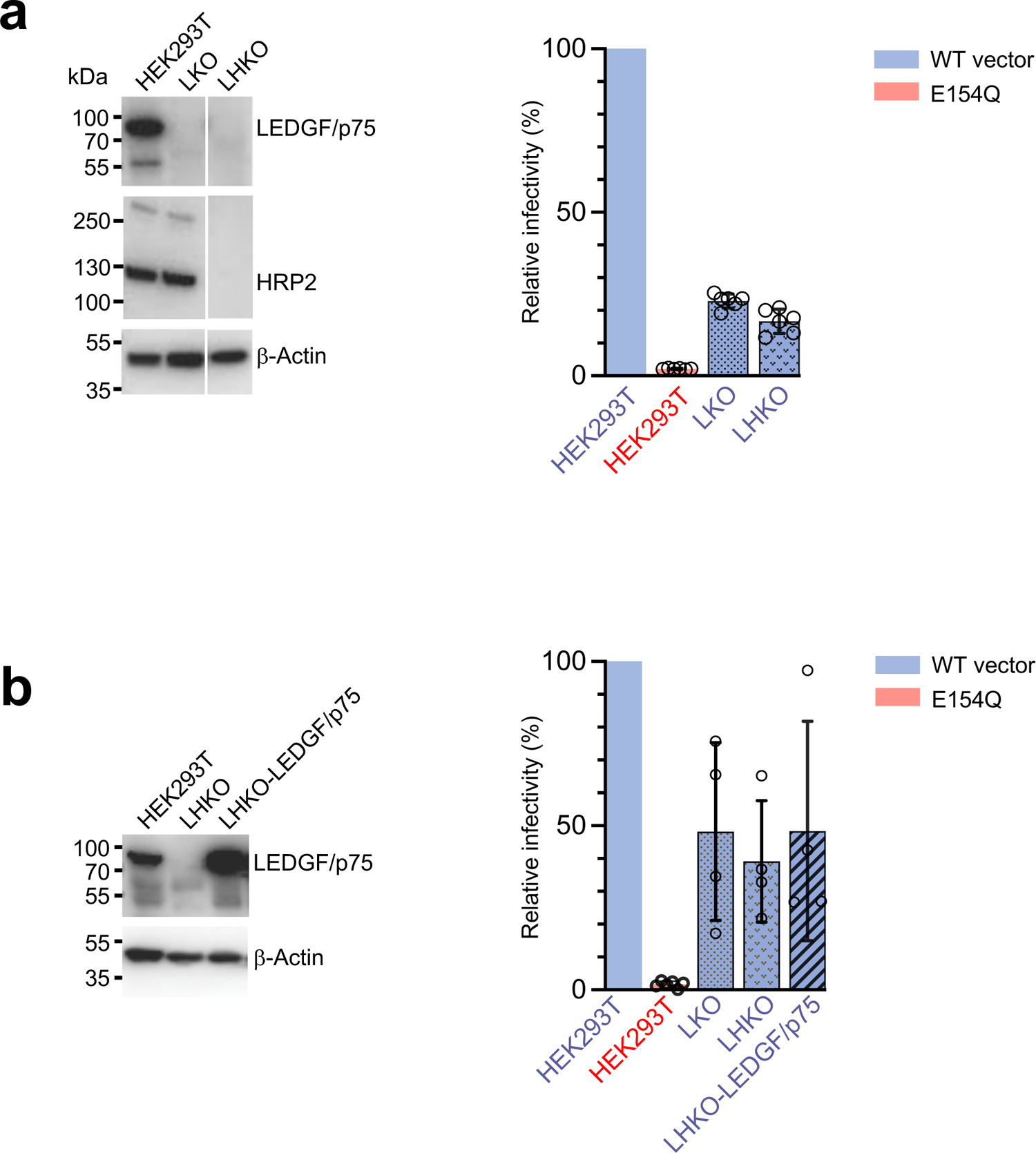
MVV vector infection of gene-modified human cells. **(A)** Infectivity of MVV vector in human LKO and LHKO cells. Left: Western blotting analysis of total cell extracts of LKO, LHKO, and parental HEK293T cells with anti-LEDGF/p75 (top), anti-HRP2 (middle) and anti-β-actin antibodies (used as a loading control, bottom). Right: HEK293T, LKO and LHKO cells were infected with WT or E154Q MVV vectors, normalized by the associated RT activity and encoding a luciferase reporter. Luciferase expression was measured 7 d post-infection; bar plots represent mean values relative to WT, which was set to 100% for each replicate series. Standard deviations were calculated from six biological replicates; open circles are the individual measurements obtained in replicate experiments. **(B)** Overexpression of LEDGF/p75 in LHKO cells does not rescue MVV vector infectivity. Left: Western blot analysis of whole cell extracts of LHKO, LHKO-cells prior and after overexpression of ovine LEDGF/p75 and parental HEK293T cells with anti-LEDGF/p75 (top) and anti-β-actin antibodies (used as a loading control, bottom). Right: The cells were infected with WT or E154Q IN MVV vectors normalized by RT activity and encoding a luciferase reporter. Expression of luciferase was measured 7 d post-infection. Bar plots represent mean values relative to WT, which was set to 100% within each replicate series. Standard deviations were calculated from four biological replicates; open circles are individual measurements obtained in replicate experiments.

**Supplementary Figure S9.**
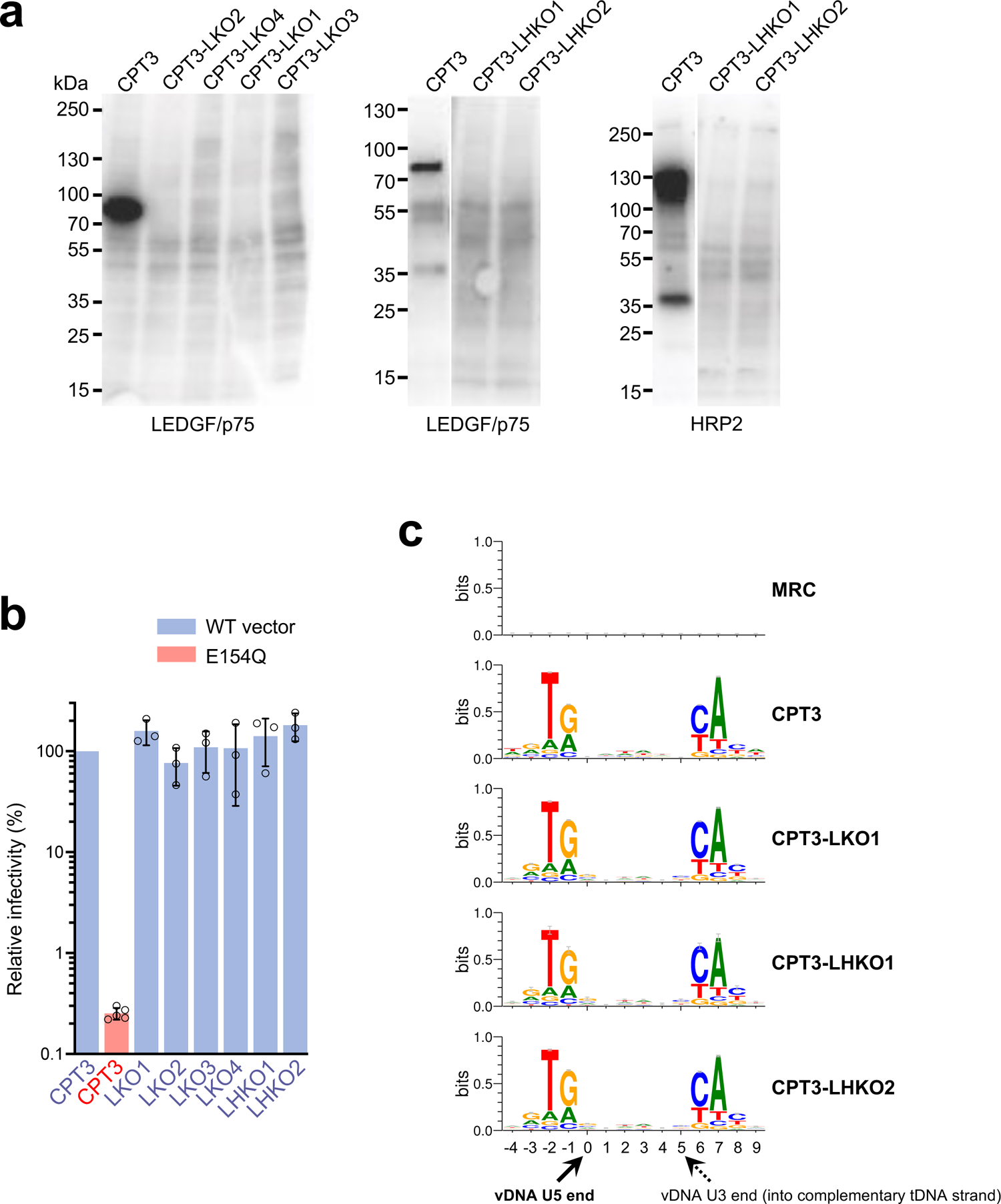
MVV infection of WT and gene-modified ovine cells. **(A)** Ablation of LEDGF/p75 and HRP2 in CPT3 cells transfected with CRISPR-Cas9 RNPs targeting ovine *PSIP1* and *HDGFL2* genes (Table S5). Equal amounts of whole cell extracts (20 μg total protein) from cells transfected with combinations of RNPs were analyzed by Western blotting with anti-LEDGF/p75 and anti-HRP2 antibodies. **(B)** MVV vector infectivity in ovine cells in the absence of LEDGF/p75 and HRP2. Ovine parental CPT3 cells or CPT3-LKO and -CPT3 LHKO cells (as indicated) were infected with luciferase-reported WT or E154Q IN MVV vectors normalized by RT activity. Luciferase expression was measured 7 d post-infection. Bars represent means; standard deviations were calculated from four replicate experiments. Open circles indicate individual replicate measurements. **(C)** Sequence logos showing local target nucleotide sequence preferences of MVV vector determined from mapped integration sites in ovine CPT3, CPT3-LKO1, CPT3-LHKO1, and CPT3-LHKO2. The mock logo (top) was generated using alignment of matched random control (MRC) sites in ovine genome.

**Supplementary Figure S10.**
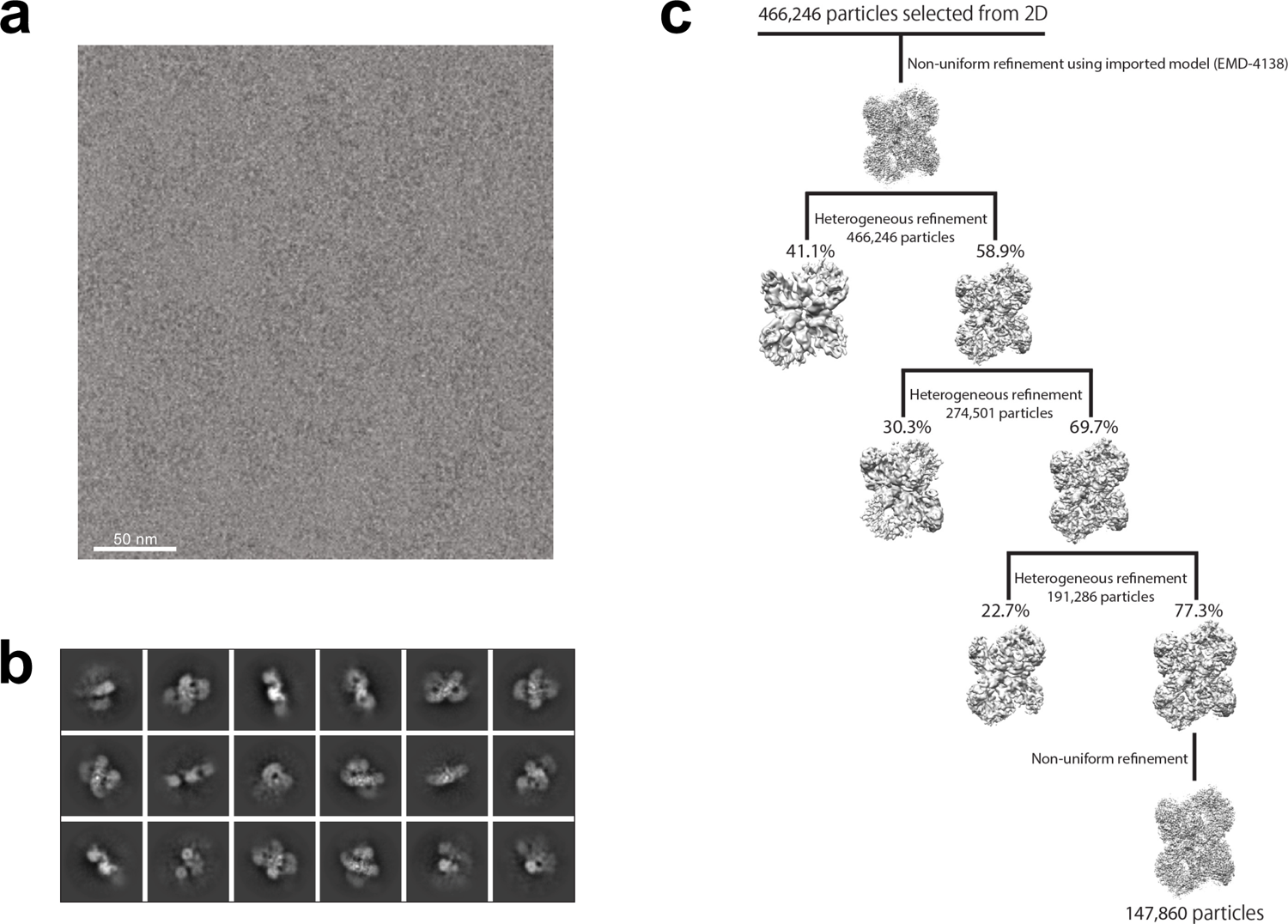
MVV CSC intasome images and classification. **(A)** Example of an image of MVV CSC particles vitrified in amorphous ice in open holes. **(B)** 2D class averages of MVV CSC particles. **(C)** 3D classification workflow utilizing iterative cycles of heterogeneous refinement and non-uniform refinement to improve the quality of the map; 147,860 particles remained for the final non-uniform refinement and reconstruction.

**Supplementary Figure S11.**
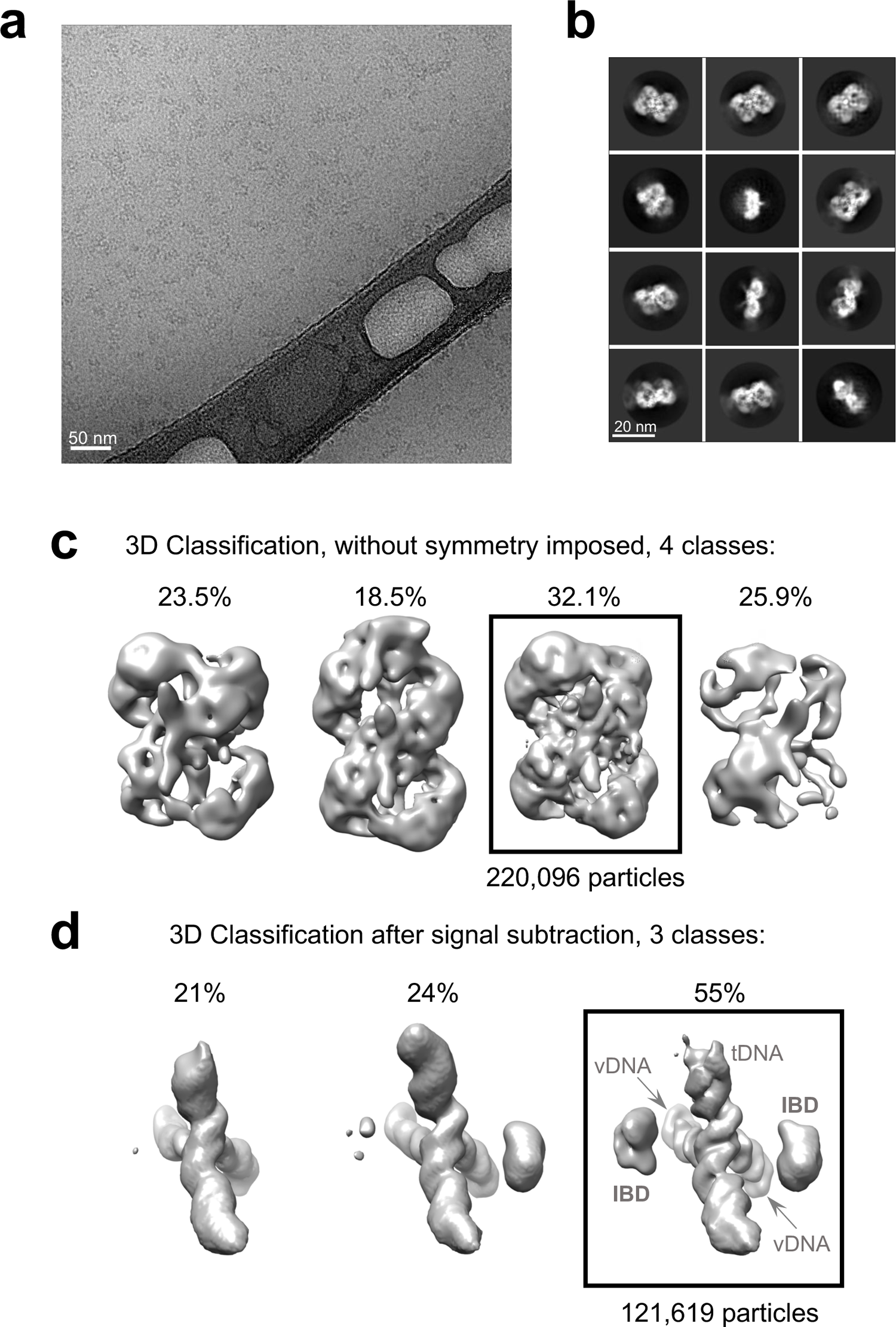
MVV STC intasome images and classification. **(A)** Example of an image of MVV STC particles vitrified in amorphous ice and supported by ultrathin carbon. **(B-C)** 2D and 3D class averages of MVV STC particles. The 3D class containing 220,096 particles (boxed in panel C) was taken for 3D classification after subtracting IN-derived signal. **(D)** Results of 3D classification following IN signal subtraction from STC particle images. The 3D class containing 121,619 particles (boxed) with good IBD occupancy at two positions was used for the final refinement and reconstruction.

**Table S1.** Cryo-EM data collection, image processing and model refinement.

|  | CSC | STC + LEDGF/p75 |
| --- | --- | --- |
| **Data collection** |  |  |
| Microscope, operating voltage | Talos Arctica, 200 keV | Titan Krios, 300 keV |
| Detector | Gatan K2 | Gatan K2 |
| Magnification (nominal) | 45,000 | 36,232 |
| Pixel size (Å) | 0.92 | 1.38 |
| Underfocus range (nominal, µm) | 1.0-3.0 | 1.5-4.5 |
| Stage tilt (°) | 40 | 0 |
| Number of frames per movie | 100 | 50 |
| Total electron fluence (e/Å<sup>2</sup>) | 43.6 | 50.4 |
| Automation software | Leginon | EPU |
| Total number of movies used | 2,295 | 11,760 |
| **Reconstruction** |  |  |
| Software for 2D classification | cryoSPARC-2 | Relion-2.1 |
| Software for 3D classification | cryoSPARC-2 | Relion-2.1 |
| Software for reconstruction | cryoSPARC-2 | cryoSPARC-2 |
| Number of initially extracted particles | 926,176 | 2,022,321 |
| Number of refined particles | 147,860 | 121,619 |
| Symmetry imposed | C2 | C2 |
| Resolution global (FSC 0.143, Å) | 3.4 | 3.5 |
| SCF value <sup>a</sup> | 0.76 | 0.84 |
| **Model refinement** |  |  |
| Software for real-space refinement | Phenix dev-4213-000 | Phenix dev-4213-000 |
| **Model composition** |  |  |
| Non-hydrogen atoms | 32,156 | 35,617 |
| Protein residues | 3,802 | 4,031 |
| Nucleic acid bases | 64 | 140 |
| Metal ions (Zn/Ca) | 12/2 | 12/0 |
| **B factors (Å<sup>2</sup>)** |  |  |
| Protein | 267.6 | 198.1 |
| Nucleic acid bases | 149.3 | 199.4 |
| Metal ions (Zn/Ca) | 148.6 | 270.72 |
| Real-space correlation coefficient | 0.70 | 0.77 |
| **R.m.s. deviations** |  |  |
| Bond lengths (Å) | 0.002 | 0.002 |
| Bond angles (°) | 0.507 | 0.51 |
| **Validation<sup>c</sup>** |  |  |
| MolProbity score | 1.47 | 1.48 |
| Clash score | 7.76 | 7.25 |
| Poor rotamers (%) | 0 | 2 |
| **Ramachandran plot quality (%) <sup>b</sup>** |  |  |
| Favoured | 97.81 | 97.63 |
| Disallowed | 0 | 0 |

<sup>a</sup> Based on the nominal Euler angle distribution<sup>b</sup> Assessed using MolProbity (96).

**Table S2.** Descriptive statistics for LEDGF/p75-Surf649 photobleaching experiments.

|  | 0.2 M NaCl | 0.5 M NaCl | 1 M NaCl |
| --- | --- | --- | --- |
| N | 605 | 745 | 516 |
| Minimum | 1 | 1 | 1 |
| 25% Percentile | 5 | 2 | 1 |
| **Median** | **6** | **4** | **2** |
| 75% Percentile | 8 | 5 | 3 |
| Maximum | 16 | 12 | 7 |
| Mean ± Std. Error | 6.41 ± 0.10 | 3.78 ± 0.07 | 2.39 ± 0.06 |
| Std. Dev. ± Std. Error | 2.45 ± 0.10 | 1.79 ± 0.07 | 1.36 ± 0.06 |
| **Intasome: LEDGF binding stoichiometry** | **1 : 6** | **1 : 4** | **1 : 2** |

**Table S3.** Statistical significance tests of HIV-1 and MVV integration site distributions in human cells.

|  | MRC | HIV-1 in HEK293T | HIV-1 in LKO | MVV in HEK293T | MVV in LKO |
| --- | --- | --- | --- | --- | --- |
| Integration in TUs <sup>a</sup> |  |  |  |  |  |
| HIV-1 in HEK293T | <10<sup>-300</sup> |  |  |  |  |
| HIV-1 in LKO | <10<sup>-300</sup> | <10<sup>-300</sup> |  |  |  |
| MVV in HEK293T | <10<sup>-300</sup> | <10<sup>-300</sup> | 2·10<sup>-40</sup> |  |  |
| MVV in LKO | 0.01 | <10<sup>-300</sup> | 10<sup>-45</sup> | 10<sup>-77</sup> |  |
| MVV in LHKO | 5·10<sup>-35</sup> | <10<sup>-300</sup> | 10<sup>-227</sup> | <10<sup>-300</sup> | 0.2 |
| Integration close TSSs <sup>a</sup> |  |  |  |  |  |
| HIV-1 in HEK293T | 3·10<sup>-5</sup> |  |  |  |  |
| HIV-1 in LKO | <10<sup>-300</sup> | 10<sup>-210</sup> |  |  |  |
| MVV in HEK293T | 10<sup>-81</sup> | 2·10<sup>-44</sup> | <10<sup>-300</sup> |  |  |
| MVV in LKO | 10<sup>-15</sup> | 2·10<sup>-11</sup> | 2·10<sup>-5</sup> | 3·10<sup>-26</sup> |  |
| MVV in LHKO | 10<sup>-159</sup> | 10<sup>-80</sup> | 5·10<sup>-22</sup> | 2·10<sup>-278</sup> | 0.7 |
| Integration near CpG islands <sup>a</sup> |  |  |  |  |  |
| HIV-1 in HEK293T | 6·10<sup>-59</sup> |  |  |  |  |
| HIV-1 in LKO | <10<sup>-300</sup> | 2·10<sup>-184</sup> |  |  |  |
| MVV in HEK293T | 10<sup>-157</sup> | 10<sup>-207</sup> | <10<sup>-300</sup> |  |  |
| MVV in LKO | 10<sup>-16</sup> | 10<sup>-4</sup> | 2·10<sup>-9</sup> | 3·10<sup>-33</sup> |  |
| MVV in LHKO | 10<sup>-223</sup> | 3·10<sup>-52</sup> | 10<sup>-29</sup> | <10<sup>-300</sup> | 0.1 |
| Integration near LADs <sup>a</sup> |  |  |  |  |  |
| HIV-1 in HEK293T | <10<sup>-300</sup> |  |  |  |  |
| HIV-1 in LKO | <10<sup>-300</sup> | 10<sup>-129</sup> |  |  |  |
| MVV in HEK293T | <10<sup>-300</sup> | <10<sup>-300</sup> | <10<sup>-300</sup> |  |  |
| MVV in LKO | 0.2 | 3·10<sup>-154</sup> | 3·10<sup>-59</sup> | 2·10<sup>-06</sup> |  |
| MVV in LHKO | 0.05 | <10<sup>-300</sup> | <10<sup>-300</sup> | 2·10<sup>-69</sup> | 0.5 |
| Integration near SPADs <sup>a</sup> |  |  |  |  |  |
| HIV-1 in HEK293T | <10<sup>-300</sup> |  |  |  |  |
| HIV-1 in LKO | <10<sup>-300</sup> | <10<sup>-300</sup> |  |  |  |
| MVV in HEK293T | 4·10<sup>-4</sup> | <10<sup>-300</sup> | <10<sup>-300</sup> |  |  |
| MVV in LKO | 10<sup>-20</sup> | 2·10<sup>-135</sup> | 10<sup>-23</sup> | 4·10<sup>-19</sup> |  |
| MVV in LHKO | 10<sup>-160</sup> | <10<sup>-300</sup> | 2·10<sup>-169</sup> | 1·10<sup>-149</sup> | 0.5 |
| Gene density near integration sites <sup>b</sup> |  |  |  |  |  |
| HIV-1 in HEK293T | <10<sup>-300</sup> |  |  |  |  |
| HIV-1 in LKO | <10<sup>-300</sup> | <10<sup>-300</sup> |  |  |  |
| MVV in HEK293T | <10<sup>-300</sup> | <10<sup>-300</sup> | <10<sup>-300</sup> |  |  |
| MVV in LKO | 10<sup>-12</sup> | 2·10<sup>-289</sup> | 10<sup>-73</sup> | 0.3 |  |
| MVV in LHKO | 2·10<sup>-47</sup> | <10<sup>-300</sup> | <10<sup>-300</sup> | 4·10<sup>-8</sup> | 0.02 |
| GC content near integration sites <sup>b</sup> |  |  |  |  |  |
| HIV-1 in HEK293T | 6·10<sup>-121</sup> |  |  |  |  |
| HIV-1 in LKO | 6·10<sup>-153</sup> | 8·10<sup>-289</sup> |  |  |  |
| MVV in HEK293T | <10<sup>-300</sup> | 4·10<sup>-84</sup> | <10<sup>-300</sup> |  |  |
| MVV in LKO | 4·10<sup>-64</sup> | 10<sup>-98</sup> | 3·10<sup>-18</sup> | 10<sup>-149</sup> |  |
| MVV in LHKO | 3·10<sup>-154</sup> | 10<sup>-282</sup> | 0.07 | <10<sup>-300</sup> | 2·10<sup>-14</sup> |

<sup>a</sup> P values determined using Fisher's exact test.
<sup>b</sup> P values determined using Wilcoxon rank-sum test.

**Table S4.** Statistical significance tests of MVV integration site distributions in ovine cells.

|  | MRC | CPT3 | CPT3-LKO1 | CPT3-LHKO1 |
| --- | --- | --- | --- | --- |
| Integration into TUs <sup>a</sup> |  |  |  |  |
| CPT3 | $<10^{-300}$ | | | |
| CPT3-LKO1 | $<10^{-300}$ | $<10^{-300}$ | | |
| CPT3-LHKO1 | $10^{-7}$ | $3 \cdot 10^{-221}$ | $3 \cdot 10^{-7}$ | |
| CPT3-LHKO2 | $<10^{-300}$ | $<10^{-300}$ | 0.2 | $2 \cdot 10^{-6}$ |
| Integration near TSS <sup>a</sup> |  |  |  |  |
| CPT3 | $10^{-24}$ | | | |
| CPT3-LKO1 | $<10^{-300}$ | $<10^{-300}$ | | |
| CPT3-LHKO1 | $2 \cdot 10^{-55}$ | $2 \cdot 10^{-43}$ | 0.004 | |
| CPT3-LHKO2 | $<10^{-300}$ | $<10^{-300}$ | 0.7 | 0.003 |
| Integration near CpG islands: <sup>a</sup> |  |  |  |  |
| CPT3 | $2 \cdot 10^{-181}$ | | | |
| CPT3-LKO1 | $<10^{-300}$ | $<10^{-300}$ | | |
| CPT3-LHKO1 | $10^{-84}$ | $2 \cdot 10^{-139}$ | 0.008 | |
| CPT3-LHKO2 | $<10^{-300}$ | $<10^{-300}$ | 0.002 | 0.07 |
| Gene density near integration sites <sup>b</sup> |  |  |  |  |
| CPT3 | $<10^{-300}$ | | | |
| CPT3-LKO1 | $<10^{-300}$ | $<10^{-300}$ | | |
| CPT3-LHKO1 | $<10^{-300}$ | $10^{-4}$ | $2 \cdot 10^{-4}$ | |
| CPT3-LHKO2 | $<10^{-300}$ | $2 \cdot 10^{-78}$ | $5 \cdot 10^{-32}$ | 0.6 |
| GC content near integration sites <sup>b</sup> |  |  |  |  |
| CPT3 | $<10^{-300}$ | | | |
| CPT3-LKO1 | $<10^{-300}$ | $<10^{-300}$ | | |
| CPT3-LHKO2 | $6 \cdot 10^{-274}$ | $<10^{-300}$ | $<10^{-300}$ | |

<sup>a</sup> P values determined using Fisher's exact test.<sup>b</sup> P values determined using Wilcoxon rank-sum test.

**Table S5.**
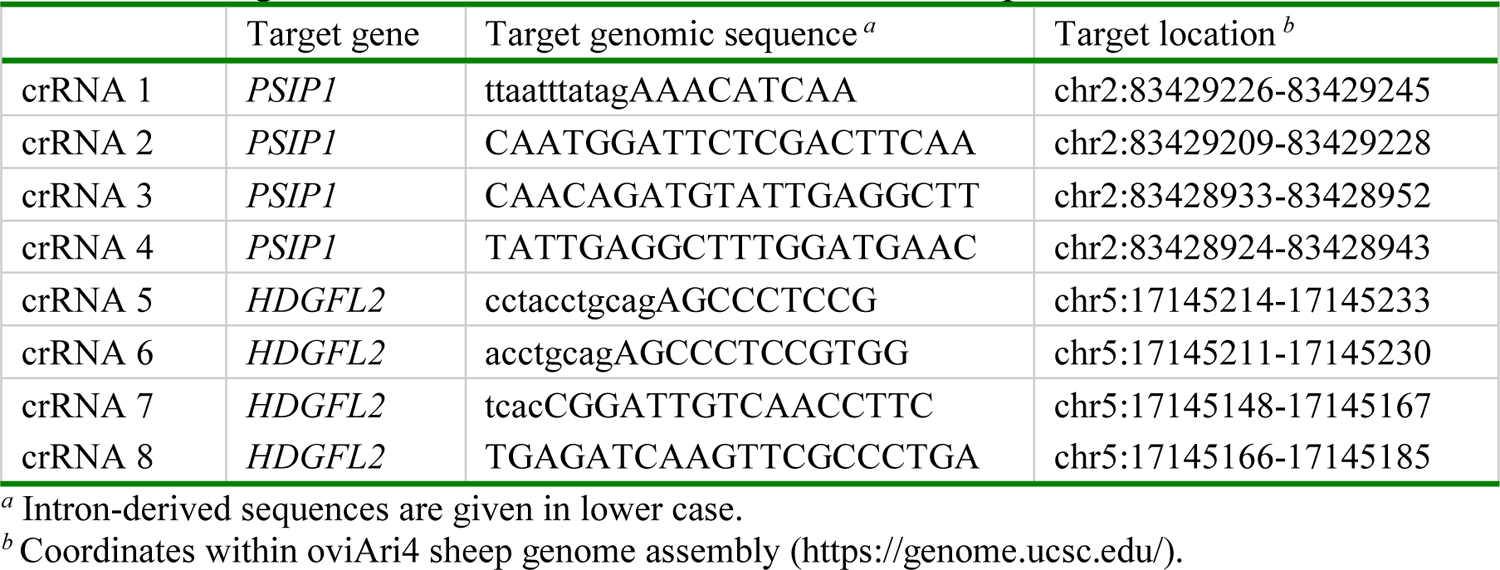
Oligoribonucleotides used as crRNAs to disrupt ovine *PSIP1* and *HDGFL2*.

